# Acquisition and processing methods of whole-brain layer-fMRI VASO and BOLD: The Kenshu dataset

**DOI:** 10.1101/2022.08.19.504502

**Authors:** Kenshu Koiso, Anna K Müller, Kazuaki Akamatsu, Sebastian Dresbach, Christopher J Wiggins, Omer Faruk Gulban, Rainer Goebel, Yoichi Miyawaki, Benedikt A Poser, Laurentius Huber

**Affiliations:** Faculty of Psychology and Neuroscience, Maastricht University, Maastricht, NL; Graduate School of Informatics and Engineering, The University of Electro-Communications, Tokyo, Japan; Imaging Core Facility, Forschungszentrum Jülich, Jülich, Germany; Brain Innovation, Maastricht, NL; Center for Neuroscience and Biomedical Engineering, The University of Electro-Communications, Tokyo, Japan

## Abstract

Cortical depth-dependent functional magnetic resonance image (fMRI), also known as layer-fMRI, has the potential to capture directional neural information flow of brain computations within and across large-scale cortical brain networks. E.g., layer-fMRI can differentiate feedforward and feedback cortical input in hierarchically organized brain networks.

Recent advancements in 3D-EPI sampling approaches and MR contrast generation strategies have allowed proof-of-principle studies showing that layer-fMRI can provide sufficient data quality for capturing laminar changes in functional connectivity. These studies have however not shown how reliable the signal is and how repeatable the respective results are. It is especially unclear whether whole-brain layer-fMRI functional connectivity protocols are widely applicable across common neuroscience-driven analysis approaches. Moreover, there are no established preprocessing fMRI methods that are optimized to work for whole-brain layer-fMRI datasets.

In this work, we aimed to serve the field of layer-fMRI and build tools for future routine whole-brain layer-fMRI in application-based neuroscience research. We have developed publicly available sequences, acquisition protocols, and processing pipelines for whole-brain layer-fMRI. These protocols are validated across 60 hours of scanning in nine participants. Specifically, we identified and exploited methodological advancements for maximizing tSNR efficiency and test-retest reliability.

We are sharing an extensive multi-modal whole-brain layer-fMRI dataset (20 scan hours of movie-watching in a single participant) for the purpose of benchmarking future method developments: The Kenshu dataset. With this dataset, we are also exemplifying the usefulness of whole brain layer-fMRI for commonly applied analysis approaches in modern cognitive neuroscience fMRI studies. This includes connectivity analyses, representational similarity matrix estimations, general linear model analyses, principal component analysis clustering, etc. We believe that this work paves the road for future routine measurements of directional functional connectivity across the entire brain.

**Graphical Abstract:** 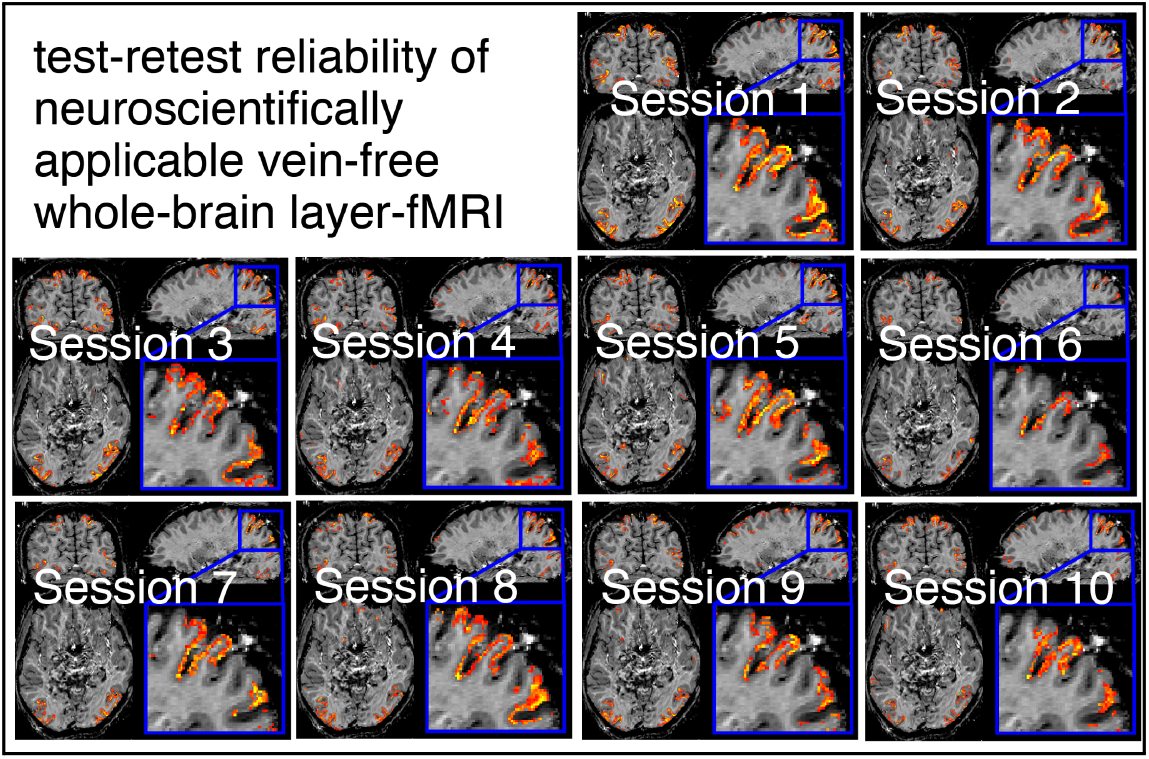

## Introduction

Cortical depth-dependent functional magnetic resonance image (fMRI), also known as layer-fMRI, allows neuroscientists to address questions of directional functional connectivity across cortical brain areas. Because of its potential to study causal neural connectivity, human layer-fMRI has become an emerging sub-field of the neuroimaging community, with approximately 40-60 yearly published papers, and doubling approximately every 2.5 years (https://layerfmri.com/papers/).

Neuroimaging with fMRI can non-invasively investigate how the brain works by means of tracking temporal activity changes within gray matter (GM). Specifically, in layer-fMRI, activation changes are captured across cortical depths and then interpreted in the context of expected neural microcircuits and neural connections terminating in specific cortical layers (Weldon and Olman 2021).

Currently, most layer-fMRI studies in humans employ imaging protocols with 0.8 mm isotropic resolutions, restricted imaging coverages (<few cm slab thickness), and temporal resolutions (TR) of several seconds (Bollmann and Barth 2021). Data are then commonly acquired for 1-4 task conditions.

### Specifically

- **Locally optimized brain coverage:** Most layer-fMRI studies usually focus on locally confined brain areas and investigate how the layer-specific fMRI signal is differentially modulated for different task conditions within restricted field-of-views (FOVs) (Schluppeck, Sanchez-Panchuelo, and Francis 2018). This experimental procedure and task design with small FOVs was vital to establishing the validity of the layer-fMRI methodology. However, restricted FOV’s do not allow the layer-fMRI methodology to fulfill its full potential to reveal directional neural information flow between brain areas of common brain networks. In fact, most of the established resting-state functional connectivity networks span across the entire cortex and cannot be fully captured with a restricted field of views. In this study, we aim to develop layer-fMRI imaging protocols that capture the entire cortex. This is to facilitate future laminar network analyses as a neuroscientific research tool.
- **Low-dimensional optimized tasks:** Another limiting factor of most layer-fMRI studies is that they employ low-dimensional activation tasks with very few task conditions. This can limit the efficiency of fMRI protocols (Naselaris et al. 2011). In fact, since the advent of human layer-fMRI (Menon and Goodyear 1999), the field of layer-fMRI has been mostly focused on methodological aspects of acquisition and analysis. Thus, the experimental design has been usually kept as conservative as possible and layer-fMRI has not yet become an established tool for use in whole-brain connectivity studies, which are of increasing importance to human neuroscience research. In order to make layer-fMRI modalities more attractive for neuroscientists to investigate neural information flow across large cortical brain networks, we aim to establish the feasibility of layer-fMRI large coverage protocols for providing reliable and repeatable connectivity estimates across cortical depth with high-dimensional naturalistic tasks of movie watching.

Previous high-resolution studies have convincingly shown that layer-fMRI acquisition protocols can be increased to capture large patches of the cortex (Finn, Huber, and Bandettini 2021; Bandettini, Huber, and Finn 2021; Huber et al. 2021; Yun et al. 2022; Pais-Roldán et al. 2020; Sharoh et al. 2019; Chang et al. 2022; Deshpande, Zhao, and Robinson 2022; Müller et al. 2021). Each of these articles shows that there are functional connectivity signal modulations across cortical depth that layer-fMRI can capture. In this study, we aim to further investigate the layer-specific signal of functional connectivity with a focus on its stability, repeatability, and neuroscientific applicability. In pursuing this, we specifically focus on practicability aspects of whole-brain layer-fMRI acquisition approaches. As such, whole brain layer-fMRI is constrained by the following aspects:

- **Slow TR:** Previous attempts at whole-brain connectome datasets had long volume acquisition times (TR_vol_ = 8-9 s) with correspondingly severe within-TR physiological noise artifacts. This had previously hampered straightforward neuroscientific applicability (Müller et al. 2021). Here we aim to take advantage of advanced 3D-echo-planar imaging (EPI) readout schemes (Poser et al. 2010, 2013) (but see also (Stirnberg and Stöcker 2021)) to mitigate those constraints.
- **Strong geometric distortions:** The relatively low readout bandwidth of high-resolution EPI readouts trains comes along with significant geometrical image distortions. While such distortions are somewhat contained for small FOV protocols and locally optimized B_0_-shimming, they impose serious applicability constraints in conventional preprocessing steps of the whole-brain layer-fMRI. For example, motion correction, run-to-run alignment, session-to-session alignment, and image registration between structural and functional data are hampered.
- **The draining vein artifact:**Conventional gradient echo (GE) BOLD fMRI contrasts are highly sensitive to trans-laminar and pial veins (Goense, Bohraus, and Logothetis 2016). This can challenge the interpretability of cortical depth-dependent connectivity results with respect to the underlying neurally-driven signal fluctuations. Here we aim to develop a vein-free cerebral blood volume (CBV) sensitive acquisition protocol using vascular space occupancy (VASO) (Lu et al. 2003; Hua et al. 2013). In VASO fMRI, an inversion-recovery pulse sequence design is used to selectively null out blood water magnetization at the image acquisition time, while leaving extra-vascular signals for detection. A CBV increase during neural activation is then associated with an overall MR-signal decrease, which in turn is believed to be proportional to the volume increase of nulled blood. In order to maintain the CBV-selective T_1_-weighing of VASO for long enough to sample the 3D k-space of submillimeter whole-brain protocols, we aim to exploit the Multiple Acquisitions with Global Excitation Cycling (MAGEC approach) (Huber et al. 2021; Lu et al. 2004; Scouten and Constable 2007).

Here, we describe our efforts to achieve a whole-brain layer-dependent functional acquisition and analysis protocol for connectome mapping of the entire cortex. The developed methodology is used to acquire a large openly available dataset of CBV and BOLD contrast. Overall, the purpose of this study is multifold:

A. We aim to implement, test, and validate an acquisition setup with minimal artifacts.
B. We seek to provide a ‘testbed’ for future efforts of developing and benchmarking new layer-dependent preprocessing and analysis tools.
C. We aim to provide at least 50 runs of movie-watching clips (15min each) to quantify the reliability of laminar connectivity results.
D. We seek to exemplify which type of new neuroscientific research questions will become addressable with whole brain layer-fMRI.

## Methods

### Participants

Nine healthy participants with normal (or corrected-to-normal) visual acuity were scanned. 44 scan hours of scanning were used to develop, optimize, and validate the acquisition protocols. 26 scan hours were used for the main experiment series of repeated movie watching. For the main scan series, one of the participants, a healthy 23 years-old male (an author (K.K.)) with normal visual and audio acuities was scanned 13 times (51 functional runs). All scanning procedures have been approved by the Ethics Review Committee for Psychology and Neuroscience (ERCPN) at Maastricht University (ERCPN-180_03_06_2017), following the principles expressed in the Declaration of Helsinki.

### Structural scanning

We acquired structural reference data at a SIEMENS MAGNETOM Prisma 3T scanner, equipped with a 32-channel head coil, operated by Scannexus B.V. (Maastricht, The Netherlands). We used an MPRAGE sequence to obtain images with a T_1_-weighted contrast for the purpose of tissue type segmentation. Protocol parameters were: echo time (TE) =2.22ms, TR=2.4s, resolution =0.8mm isotropic, flip angle =8°, field of view = 256×256×256mm. The motivation for collecting data at 3T (as opposed to 7T) was to obtain types of tissue contrast that the major processing software packages like FreeSurfer have been optimized for. Furthermore, 3T structural data can be less constrained by spatial intensity inhomogeneities, while maintaining an acceptable contrast-to-noise-ratio (CNR) (Polimeni et al. 2018). To increase the CNR of 3T structural scans, we acquired a collection of 10 averages. We were inspired to use the procedure of obtaining multiple structural references from a different scanner (3T) than the functional experiments (7T) by (Allen et al. 2022).

### Functional scanning

High-resolution functional data were obtained on a SIEMENS classic MAGNETOM 7T scanner, equipped with a 1Tx / 32Rx head coil (Nova Medical, Wilmington, MA, USA) operated by Scannexus B.V. (Maastricht, The Netherlands). For layer-fMRI scanning without venous biases, a blood volume sensitive VASO (Lu et al. 2003) 7T-sequence (Hua et al. 2013) was used. We used the MAGEC VASO approach to maintain the VASO T_1_-weighting across long echo trains (Huber et al. 2021). One 18×18×0.5cm^3^ high-permittivity dielectric pad containing a 2.8:1 solution of calcium titanate (CaTiO_3_) and heavy water (D_2_O) by weight was placed on the right side of the participants’ heads at the level of the temporal lobes to increase B_1_^+^ efficiency at 7T (Webb 2011). A 3rd-order B_0_-shim protocol was performed with three iterations using vendor-provided tools and in-house developed scripts executed at the scanner console.

### Pilot experiments on optimizing CBV contrast and in-plane readout

15 two-hour sessions in nine participants (including 2 two-hour sessions in the main participant) were performed to optimize the blood volume contrast preparation and in-plane EPI readouts. All those experiment series were conducted with 0.8mm resolution, partial Fourier=6/8, whole-brain coverage, and TE=18ms. Specifically:

- We investigated the benefit of MAGEC VASO compared to a traditional SS-SI VASO approach. We found higher tSNR values for traditional compared to MAGEC VASO. However, the T_1_ contrast-to-noise ratio and the functional CBV CNR (for flickering checkerboard stimuli) were higher for MAGEC VASO. Thus, we decided to use MAGEC VASO for the remaining experiments of the study.
- We investigated what regime of excitation flip angles would provide an optimal VASO contrast. We tested nominal flip angles of 6°, 9°, 12°, and variable flip angles ranging from 22.1° to 40° (for more information about the purpose of using variable flip angles in VASO see (Huber et al. 2018)). We found a favorable compromise between T_1_ CNR and BOLD tSNR for an interleaved approach. In all later experiments (including the main experiment) we used alternating flip angle schemes for VASO and BOLD volumes. For VASO, we used variable flip angles (22.1° - 40°). For BOLD, we used constant flip angles of 9°.
- We investigated the impact of 3^rd^-order vs. conventional 2^nd^-order shimming for whole-brain 3D-EPI readouts. We found that the improved artifact level of the 3rd-order shimming justifies the extra time investment of 4-6 minutes of higher-order shimming with three iterations.

The results of these pilot optimization experiments, which are not the focus of this article, are described in the scientific report (Müller 2020). fMRI data of these pilot scans are included in the public data repository of this study.

### Piloting experiments on optimizing parallel imaging and sampling speed

In 6 participants, we tested a series of sampling approaches across a wide spectrum of volume acquisitions durations and k-space trajectories. Specifically, we focussed on acceleration factors (AF) and Controlled Aliasing in Volumetric Parallel Imaging (CAIPI) FOV shifting (Breuer et al. 2006). The choices of the tested protocols were inspired by previous large coverage 3D-EPI studies (van Mourik et al. 2021; Poser et al. 2013; Stirnberg and Stöcker 2021):

1. AF=3 (3×1), TR=9.8s, Fat saturation per shot, no IcePat (no CAIPI).
2. AF=6 (3×2), TR=5.1s, Fat saturation per shot, CAIPI: y shift 3 (k_y_-prephased CAIPI a.k.a. shot-selective)
3. AF=9 (3×3), TR=4.8s, Fatsat: per shot, CAIPI: y shift 3 (k_y_-prephased CAIPI a.k.a. shot-selective)
4. AF=8 (4×2), TR=4.1s, Fatsat: none, CAIPI: y shift 1 (k_y_-prephased CAIPI a.k.a. shot-selective)
5. AF=6 (3×2), TR=5.1s, Fatsat: none, CAIPI: y shift 3 (k_y_-prephased CAIPI a.k.a. shot-selective)
6. AF=6 (3×2), TR=4.9s, Fatsat: none, CAIPI: z shift 2 (k_z_-blipped CAIPI)
7. AF=12 (6×2), TR=2.7s, Fatsat: none, CAIPI: y shift 3 (k_y_-prephased CAIPI a.k.a. shot-selective)
8. AF=16 (4×4), TR=2.3s, Fatsat: none, CAIPI: z shift 2 (k_z_-blipped CAIPI)

The functional usability of each of those protocols was further tested with a visuomotor task in two participants (single session each). Furthermore, the two most promising protocols (option #1 and #5) were tested with movie-watching tasks in two participants (5 sessions with 5 runs, each). All protocols had whole-brain coverage and MAGEC VASO enabled. Based on the tSNR-efficiency, artifact levels, and activation map quality (Fig. 3-4), we chose option #5 for all subsequent movie-watching experiments. This is: AF=6 (k_y_=3×k_z_=2), TR=5.1s, no Fatsat, CAIPI: y-shift 3). Note that the sampling scheme of GRAPPA 6 (k_y_=3×k_z_=2) with CAIPI y-shift 3 is identical in trajectory and sampling pattern as an in-plane segmented approach of GRAPPA 6 (k_y_=6×k_z_=1), with segmentation factor 2 and CAIPI z-shift 1. Representative results of these experiment series are shown in Fig. 3 and 4.

### Detailed protocol parameters of the main movie-watching experiments

The acquisition protocol parameters that we used for the 51 runs of movie watching were: resolution=0.84mm iso, matrix size=225×225, number of slices=112, TE=18ms, TR_vol_=5.1s (alternating 5.1s/5.2s when including dead times), FOV=184.3×184.3mm, 3D-EPI (Poser et al. 2010), GRAPPA=3×2, two segments, and 3D-CAIPI 1 (equivalent to GRAPPA 6×1 with pre-phased, shot-selective CAIPI 3) (Poser et al. 2013; Stirnberg and Stöcker 2021), acquisition time = 15min, ⩾5 runs per session. The last two TRs were NORDIC (Vizioli et al. 2021) noise scans (not included in reconstruction). The MR-sequence used here is available via SIEMENS’ C2P ‘app store’ Teamplay. Complete protocols of acquisition parameters: https://layerfmri.page.link/WBprotocol. Respiratory and cardiac traces were recorded with the vendor-provided sensors and sequence-embedded data storage for potential future vascular reactivity analyses.

The readout gradients of this protocol were retrospectively measured by means of field monitoring with 16 F19 NMR probes (Skope, Zurich, Switzerland) placed around the iso-center of the scanner. This was done to quantify the deviation of the read-gradient pulses from the nominal trapezoidal shape, which informed the choice of later artifact correction methods during preprocessing.

### Reconstruction

All images were reconstructed using a vendor-implemented GRAPPA (Griswold et al. 2002) algorithm compatible with 2D CAIPIRINHA (Breuer et al. 2006) using (‘IcePAT’) (Kellman and McVeigh 2005) with a 3D GRAPPA kernel of k_x_=5, k_y_=4, k_z_=3 and using 90(k_y_)×24(k_z_) reference lines. Partial Fourier reconstruction and coil combination was performed within IcePAT. EPI phase correction was conducted with the vendor algorithm IsOnlinePCCrossCorrAcrossSegmentsEPI (Heid 1997). The reconstruction was performed on the vendors’ imaging server MRIR and the reconstruction of each 15 min run lasted about 40min. Since the scanner was not responsive during this reconstruction time, all experiments were conducted as the last session of the day after 8 pm.

### Stimuli in validation experiments

In order to implement and optimize the imaging protocol for the naturalistic tasks, we first aimed to attest to the quality of the acquisition procedure with a visuomotor ‘validation task’ of predefined expected activation patterns. During activation periods, participants were asked to tap their index finger and thumb in a pinch-like motion, while looking at a flickering checkerboard. The stimuli and rest blocks were approximately 30s each. For comparisons of protocols with different TR lengths, we matched the task timings to the closest TR of 30s. During rest periods, participants were presented with a central fixation cross. Participants were asked to look at the center of the screen at all times.

### Stimuli in the movie-watching experiment

As soon as the acquisition protocol was validated with block-designed visuomotor tasks, we started to acquire the Kenshu dataset with a movie-watching task. We chose to use naturalistic movie-watching tasks in favor over other, more controlled task environment for multiple reasons:

- In order to fully take advantage of the whole-brain coverage of the acquisition protocol here, we were aiming for an experimental setup that facilitates functional connectivity analyses. Such analyses are commonly applied with either resting-state or naturalistic tasks.
- Compared to resting-state paradigms, brain activity fluctuations induced during repeated movie-watching are more repeatable and more consistent across multiple runs. This facilitates across-run averaging, which can be advantageous in the thermal-noise limited regime of sub-millimeter fMRI. Furthermore, this also facilitates test-retest analyses quantifying the feasibility of the developed acquisition protocols to address neuroscience research questions that commonly require relatively high robustness.
- Movie-watching paradigms are known to engage all brain areas to some degree. This facilitates exploring the quality of the developed acquisition and analysis protocols beyond isolated individual brain areas. This is important for judging the generalizability of previous lab-based layer-fMRI connectivity findings (Huber et al. 2021).
- Movie watching tasks engage the brain more than resting-state paradigms and result in larger effect sizes of fMRI signal changes (Finn and Bandettini 2021). This is particularly helpful for layer-fMRI, which is inherently constrained by detection sensitivity limitations.
- fMRI protocols with movie-watching paradigms usually have reduced head motion compared to resting-state paradigms (Vanderwal et al. 2015). This mitigates one of the challenges of layer-fMR; voxel-displacements.

We used the 15 min (180 TR_vol_) collection of movie clips that are established for exploring advanced fMRI methodology from the 7T HCP study (a.k.a. MOVIE1). This collection consists of 5 separate short stories (1:03-4:05 min) interspaced with 20 s of rest (indicated by the word “REST” in white text on a black background). The clips are from independent films freely available under a Creative Commons license and are entitled: Two Men (2009, http://vimeo.com/17970306), Welcome to 221 Bridgeville (2011, http://vimeo.com/31318354), Pockets (2008, http://vimeo.com/14216866), Inside the Human Body (2011, http://vimeo.com/24930096), 23 Degrees South (2011, vimeo repeat), and LXIV (2011, vimeo repeat). Movie-watching sessions were conducted over the course of 10 consecutively weekly sessions. No session lasted longer than 2h. In each session, the participant underwent 5 viewings (runs) containing all 5 movie clips, each. In the 9th session, one additional 6th run was conducted.

No gaze fixation was applied during movie watching. This is in accordance with the HCP study that had previously established these movie clips as an fMRI task and can be taken as a low-resolution reference dataset. Not enforcing visual gaze fixation has the advantage that it is a more ‘natural’ form of naturalistic stimulus presentation and, thus it is less taxing for the participant to undergo more than 50 movie viewings. The disadvantage of refraining from a fixation task is that some aspects of retinotopically specific stimulus processing of the movie is not fully consistent across runs. However, based on previous research of repeated movie-watching, we believe that the gaze trajectory is very consistent across runs. Namely, eye tracking data from repeated experimental runs by (Mandelkow, de Zwart, and Duyn 2017) showed very similar gaze trajectories with inter-experimental cross-correlation coefficients above 40% on average. The remaining stimulus-driven neural fluctuations that are not consistent across runs and will be treated as being part of the class of spontaneous neural activity that is anyway continuously ongoing besides the task.

Audio sounds of the movie clips were presented to the subjects in the MRI scanner using the MRI-compatible earbuds of Sensimetrics Corporation (www.sens.com).

### Data pre-processing

Fig. 2 depicts a schematic overview of the pre-processing pipeline of this study. Each step was adjusted and optimized for the specific requirements and challenges of the whole-brain layer-fMRI. Exported images from the scanner were converted from DICOM files to NIFTI files in dcm2niix (version v1.0.20210317) (Li et al. 2016). Retrospective motion correction was performed as a nonlinear alignment of each time frame to a template from the very first session using rigid, affine, and non-linear SyN algorithms in ANTs (version 2.1.0) (Avants et al. 2014). This was separately conducted for images with and without blood nulling. BOLD correction of VASO was performed in LayNii (version 2.2.0 (Huber et al. 2021)). We found that non-linear motion correction was necessary to account for head motion across areas suffering from local geometric distortions (Müller and Huber 2020). This was in contrast to previous slab-selective imaging protocols of this sequence (Huber, Finn, et al. 2020; Huber, Poser, et al. 2020), where 3^rd^-order shimming could sufficiently minimize nonlinear distortions across small motion displacements. For quality comparisons of the linear and non-linear motion correction strategies and the necessity to use such computationally expensive preprocessing procedures, see (Müller and Huber 2020). In order to mitigate signal blurring due to spatial interpolation, volume-to-volume motion correction, alignment across runs, and alignment across sessions were applied in one single interpolation step with a spline interpolation function. Structural T_1_-weighted images from 3T were aligned in ANTs and averaged (N=10 images). This was done to maximize GM/WM/CSF contrast for segmentation (Allen et al. 2022).

**Fig 1.**
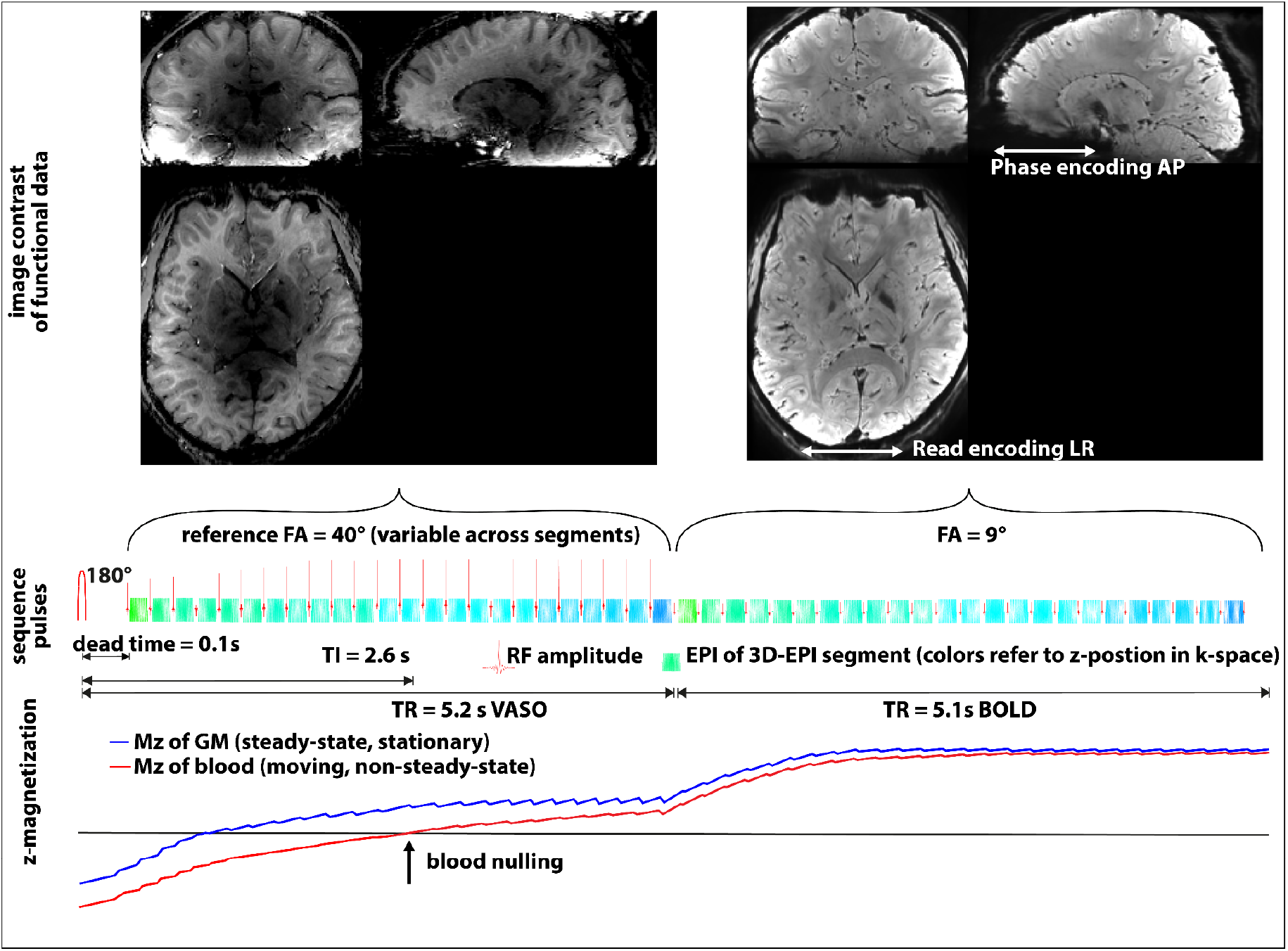
Schematic sequence diagram and expected z-magnetization of the optimized protocol used here. In SS-SI VASO with MAGEC, two contrasts are acquired concomitantly, VASO and BOLD. The acquisition of each pair of images starts with an adiabatic inversion pulse. This imposes a T_1_-contrast in the stationary brain tissue and nulled once-inverted (non-steady-state) blood magnetization at the k-space center of the subsequently acquired VASO image. In order to maintain the blood-volume sensitive T_1_-contrast for relatively long readout durations, we employed the MAGEC approach and used large excitation flip angles that prohibit a free relaxation back to equilibrium. The acquisition of the second volume (BOLD control) is acquired with a smaller flip angle and without a preceding inversion pulse. This results in a not-nulled conventional BOLD contrast. Flip angles refer to nominal values and are subject to spatially varying B ^+^ inhomogeneities at 7T. The depicted TI and TR values shown here refer to the protocol #5, which was used for the main experiment series.

**Fig 2:**
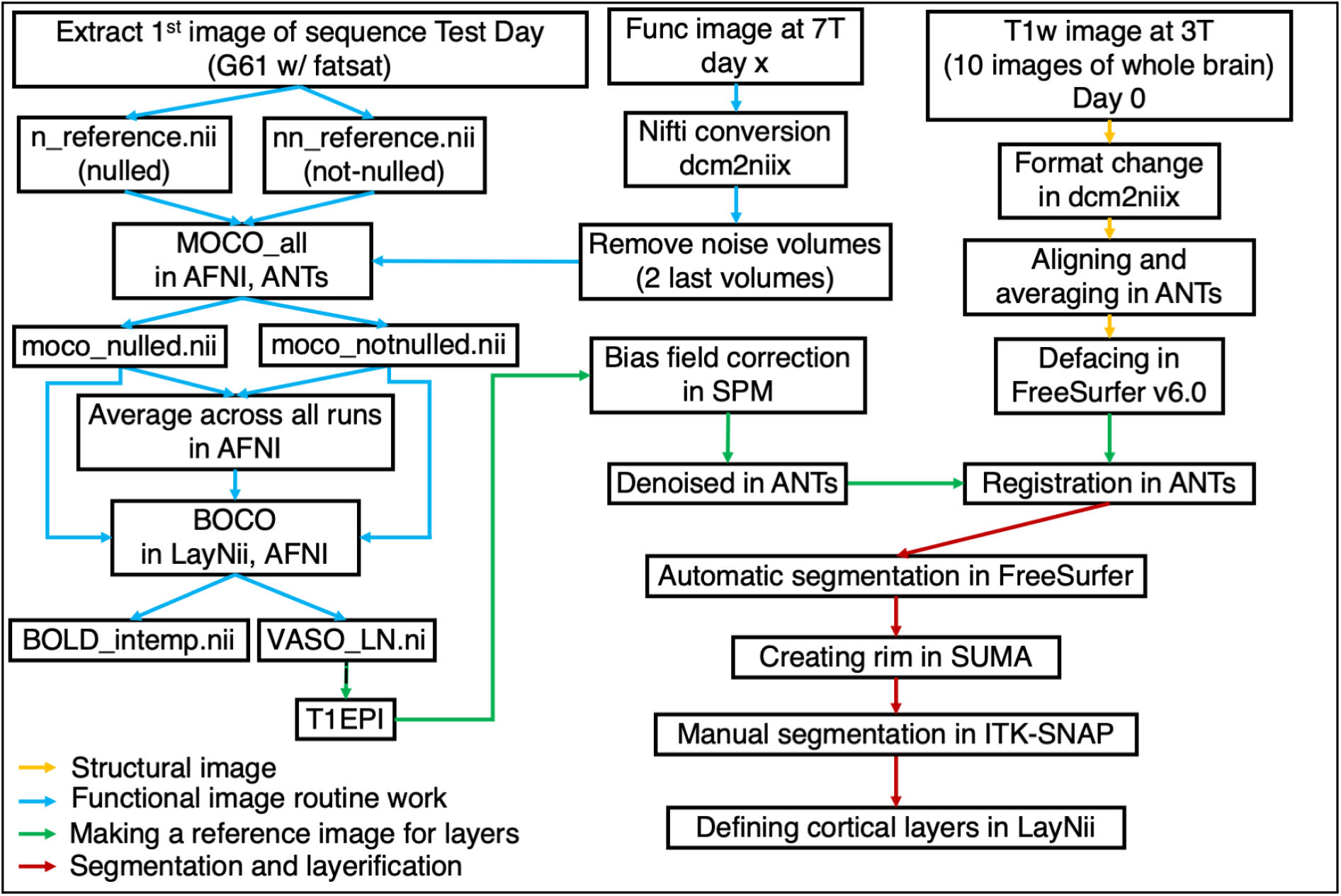
Pre-processing pipeline workflow of whole-brain layer-fMRI data While standardized and vetted fMRI preprocessing pipelines exist, they are not optimized for, and are not suitable for whole-brain layer-fMRI data. As such, layer-fMRI suffers from more severe temporal varying geometric distortions, while having higher localization quality requirements. Thus, we developed, implemented, and validated an analysis pipeline for whole-brain layer-fMRI. A schematic representation of the final preprocessing analysis flow and software is shown in this figure. The pipeline entails: structural image processing, functional image processing, and segmentation and layerlification. Each script and its order in the analysis workflow can be found on GitHub (https://github.com/kenshukoiso/Whole_Brain_Project).

Face-removal anonymization of the structural data was performed before data publication on OpenNeuro using the FreeSurfer command mri_deface_osx (version March 2020 https://surfer.nmr.mgh.harvard.edu/ftp/dist/mri_deface/).

As an EPI reference image for later layerification, we used the run-average T_1_-weighted EPI scan (Grappa 6 (3×2) with fat saturation) from the sequence comparison sessions (No. 1). These data were B_1_ bias field corrected in SPM (version 12) and denoised with a diffusion-denoising filter in ANTs. Note that this spatial filter was applied to guide alignment and segmentation. This filter was not applied for any of the functional results presented here. Finally, we registered the structural image to the functional images using SyN in ANTs.

Initial CSF/GM/WM segmentation was performed in Freesurfer (version 7.2.0). Since the segmentation was not good enough for layerification yet, we manually corrected the WM file and performed the topological correction in FreeSurfer (using WhiteMatterEdits_tktools and TopologicalDefect_tktools). Because the segmentation was still not acceptable for layer-fMRI research questions, we performed further manual correction in ITK-SNAP (version 3.8.0 (Yushkevich et al. 2006)) and a 16-inch WACOM drawing display (Kazo, Saitama, Japan). During manual segmentation corrections we focused on residual segmentation imperfections from the FreeSurfer output, including:

- WM misdetection errors that were re-introduced by FreeSurfers topology defect correction upon first manual edits. This was particularly important for parts of the calcarine sulcus.
- In some instances, parts of the dura mater and part of large venous sinuses were wrongly segmented as GM.
- Residual alignment errors between the structural and functional data in frontal brain areas resulted in mislocated GM segmentations.

For accurate manual correction, the GM segmented file (a.k.a. rim file) was created in SUMA (AFNI) with a higher resolution (upsampled to 0.4 mm isotropic). For translation of surfaces to volume space, a vertex density of 20,000,000 per hemisphere was used. In order to mitigate ‘kissing gyrus’ artifacts, one iteration of erosion and dilation was performed (for details see 3dmask_tool command in s07_MC2Layering.sh in the link below). Then, we manually corrected the segmentation file (rim file), which was done by an author (KK) and took more than 60 hours of manual labor. To further ensure the quality of the resulting segmentation, it was manually inspected and re-iterated by two experts (LH, OFG). Layerification was done in LayNii with equi-volume sampling for 11 layer-groups and prepared for usage at the original fMRI resolution (0.8 mm isotropic). For further layer analysis, we merged these 11 layer-groups into three layer-groups.

Lastly, image orientation nii headers were adapted such that the layerification results that are derived from SUMA refer to be in the same reference space as the native functional data. This was also applied to brain area reference atlases (Glasser 2016) to use them in the EPI reference space later.

All preprocessing scripts are openly accessible here: https://github.com/kenshukoiso/Whole_Brain_Project

### Layer-connectivity analysis

To investigate connectivity networks across layers, we performed independent component analysis (ICA) on run-averaged functional time series across different layer bins. For simplicity, we used only three independent layer bins. We applied ICA in FSL MELODIC (version 6.0.5.1) with 30 components. In order to exemplify the replicability of movie networks across sessions, we manually selected one IC and extracted its time course. This time-course was then used as a regressor for session-specific general linear model (GLM) analyses conducted in FSL Feat (version 6.0.5.1).

For the columns of specific layer profiles, we defined 10000 geometrically-defined column-like ROIs (LayNii LN2_COLUMNS), calculated ‘hubness’ in AFNI’s ‘3dTcorrMap’ (version AFNI_20.3.05), and performed principal component analysis (PCA) with seven components in AFNI ‘3dpc’ (version AFNI_20.3.05) of the hubness layer-profiles (with 7 layer-bins, 5 within GM) across all columns. Then, we highlighted individual layer profiles and presented their columnar distribution across the brain. More graphical explanations about this procedure can be found in Fig. 5 of (Huber et al. 2021).

### Multivariate pattern analysis and temporal similarity analysis

For representational similarity analyses (RSAs), we first averaged all 51 runs to maximize the functional SNR. For additional processing, detrending in AFNI ‘3dDetrend’ (command with 3^rd^-order polynomials) was applied to remove scanner-induced signal drifts. We then aimed to quantify the temporal evolution of layer-dependent representations of each movie frame and calculated the representational dissimilarity matrix (RDM) (Kriegeskorte 2008) volume-by-volume in each brain area in three independent layer groups, respectively. Brain areas were defined by the Desikan-Killiany-Tourville (DKT) atlas (Klein and Tourville 2012) and the Glasser atlas (Glasser et al. 2016).

In order to capture variations of similarity values across movie structure (movie clips and rest periods) more clearly, dissimilarity values within each movie clip were averaged. Also, to depict the layer-specific features of the similarity across brain areas, we calculated the average within the half area of matrices in each three-layer group and each brain area. Schematic depictions of these analysis steps are shown in Fig. 9A (and https://youtu.be/BAIMgMr0Ygg?t=619).

In order to validate that the representational similarly results are not biased and not driven by artifacts in the acquisition procedure (e.g. residual scanner drifts, frequency instabilities), we replicated the analysis with a pair of datasets. Namely, we first separated our data by odd and even sessions and then averaged them respectively. Ultimately, we calculated RDMs, clip-averaged RDMs, and similarity layer profiles in the same way as described above. In this analysis, the representational similarity across different movie frames is estimated on independently collected data. The results of this validation analysis are summarized in Fig. 10. This analysis was independently conducted across all brain areas in the atlas and all layers. While the results presented here focus on selected visual areas, all results can be browsed here.

In order to compare the obtained similarity analysis results to behavior data, we also performed a behavior experiment. This time, the main participant was asked to rank the similarity of pairs of movie clips from 0 to 10. Since this main participant is an author, this experiment was performed before data analysis.

In addition to the RSAs described above, we also investigated the similarity of temporal signal variations across layers and brain areas. Such temporal similarity analyses are more widely known as ‘functional connectivity’. Specifically, we extracted the spatially averaged fMRI signal traces of three layer groups in visual brain areas (V1-V8). Then, the Pearson correlation values were calculated for all combinations of layers and visual brain areas and sorted in a matrix form, a.k.a. connectivity matrix. Results of this analysis are summaries in Fig. 11.

### Estimation of vascular reactivity

Furthermore, we estimated measures of cerebral vascular reactivity (CVR) based on the amplitude of low frequency fluctuations in the BOLD signal. Specifically, we used the Physiological Basis of Vascular Autocalibration (VasA) approach (Kazan et al. 2016, 2017). In this approach, all 51 BOLD runs were transformed into temporal frequency space with FSL (fslpspec), demeaned, and normalized with the G-factor map. Then, the averaged power within the frequency band of 0.01–0.08 Hz was extracted across cortical depth for all 180 cortical areas in the Glasser atlas. This estimate of CVR is expected to be spatially highly correlated with the relative distribution of venous baseline blood volume (Kazan et al. 2017) and it is very similar to alternative metrics known under that names RSFA (Guidi et al. 2020; Liu et al. 2013) or ALFF (Chen and Gauthier 2021).

## Results

Fig. 3 shows the effect of newly incorporated and tested sequence advancements compared to the approach of simply increasing the number of slices from previous layer-fMRI protocols with restricted FOVs. The reduction of spatial and temporal artifacts with segmentation and shorter TRs are clearly visible. We believe that the different artifact level is mostly related to the different effective echo spacing. Refraining from CAIPI blips in favor of shot-selective CAIPI shifts allowed us to keep the shot-to-shot B_0_-variations in the k_z_-direction completely orthogonal to the low bandwidths in k_y_-direction. This helps to minimize intermittent ghosting artifacts across the EPI time series.

**Fig 3:**
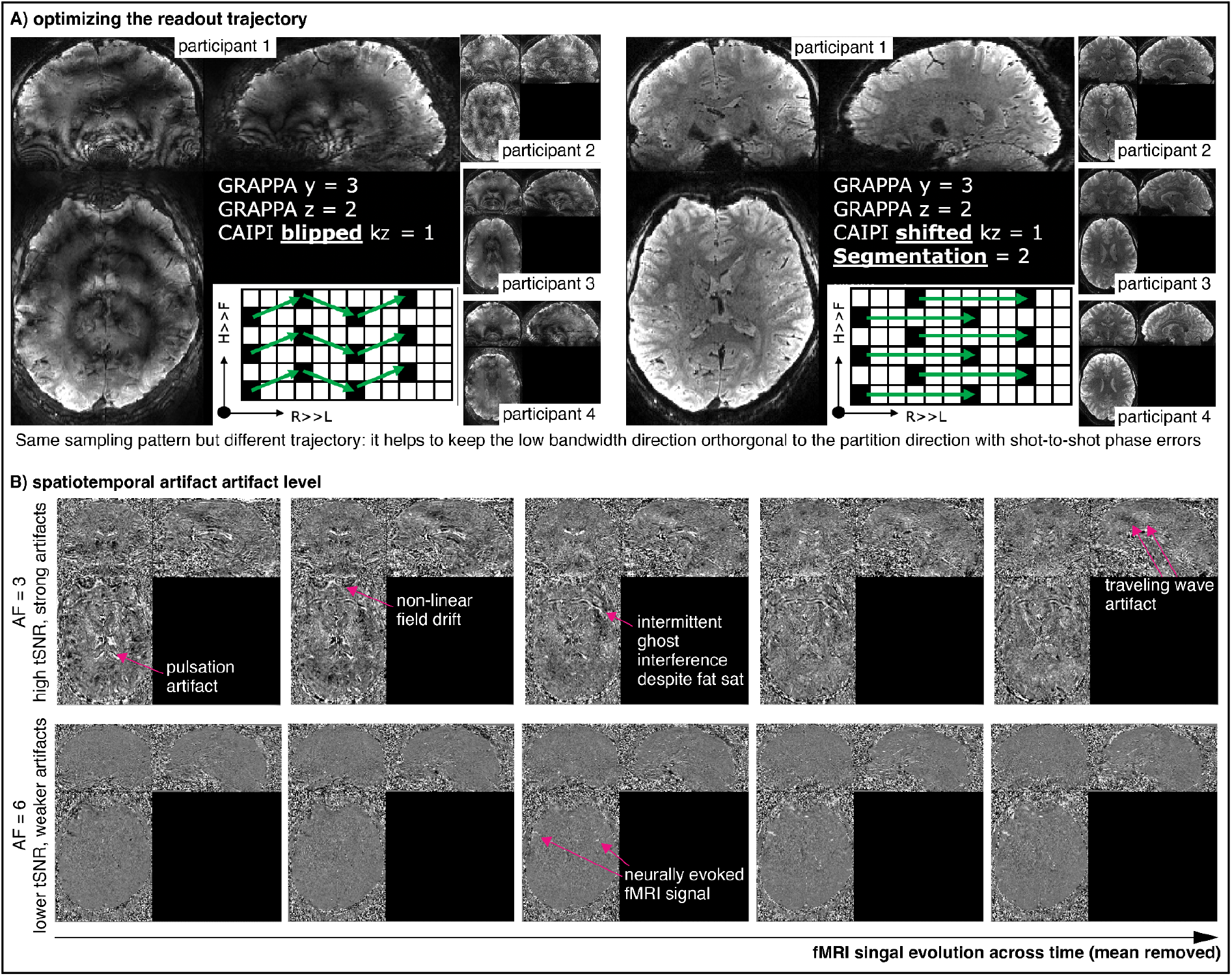
Artifact level across protocols. For an animated version of this figure, please see here: Panel A: Segmentation and k_z_-CAIPI approaches allowed us to keep artifact levels at bay and benefit from relatively liberal acceleration factors (AF=6) and relatively short TRs without prohibitive ghosting artifacts that are common when mixing the low bandwidth direction with the partition direction. This insight allowed us to reduce the volume TR from 8.9s to 5.1s. Panel B: Temporal artifact level for AF=6 and AF=3. This panel depicts the temporal VASO signal evolution (detrended) of movie watching data averaged across 25 runs each. It can be seen that shorter TRs with a shorter effective bandwidth result in improved interpretability of temporal signal variations. This is despite the overall reduced tSNR efficiency of AF=6 compared to AF=3 (see Fig. 4A).

Fig. 4 depicts representative results from pilot experiments across a large spectrum of acceleration parameters. We find that a six-fold acceleration with a k_y_-prephased (a.k.a. segmented) approach is a good compromise between physiological noise artifacts and G-factor noise amplifications. Due to the short fat T_2_^*^ at 7T, most of the fat signal appeared to have already decayed away before the TE of 18ms, and a shot-selective fat saturation did not improve the image quality enough to justify the respective time and SAR allocation. Thus, fat saturation (Fatsat) was not applied for subsequent experiments.

**Fig 4:**
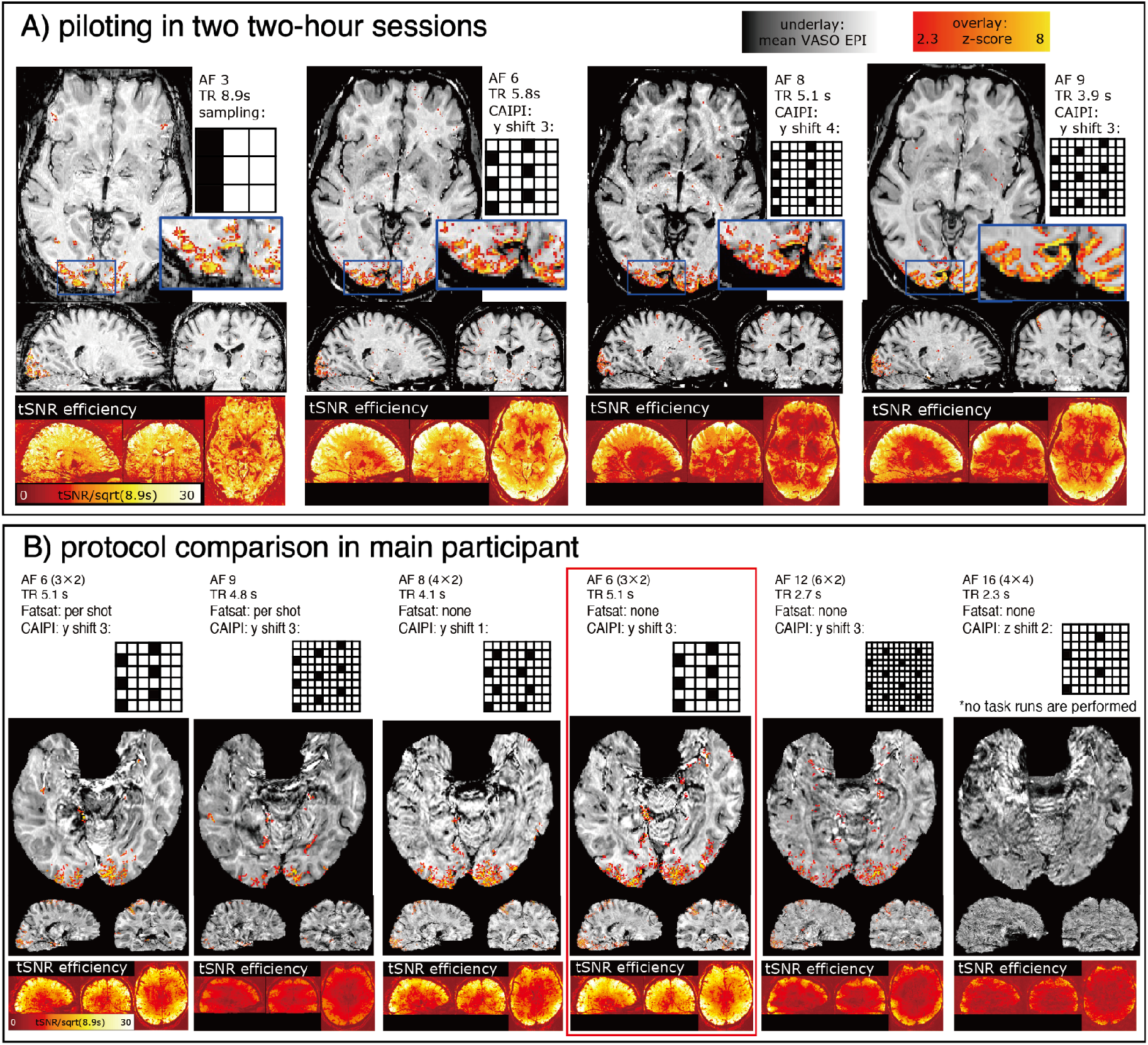
Piloting GRAPPA undersampling schemes Whole-brain sub-millimeter fMRI data contain more than 5,000,000 voxels per volume. It takes time to acquire so much data, effectively limiting the temporal resolution. In order to find a suitable compromise between minimal GRAPPA acceleration, short TR, maximal tSNR-efficiency, and acceptable artifact level, we tested multiple protocols in a series of pilot experiments. This figure shows a test series of two representative subjects. Experiments were conducted with a visuomotor task. Panels A and B refer to different participants. Panel A shows acceleration factors 6 can provide relatively high tSNR and activation maps without large losses in efficiency (TR=5.8s, CAIPI y-shift 3). Panel B depicts a series of even more liberal acceleration factors. These data are acquired in the main participant that also underwent the 51 run movie-watching task. The fourth column (AF=6, TR=5.1s, red box) depicts a good compromise of TR and artifact level. We decided to use this sequence for all subsequent scans. All activation maps refer to VASO signal changes (vein-contaminated BOLD data are not shown here).

Fig. 5 depicts the preprocessing quality including: alignment of the anatomical image to functional space, layerification in EPI space, alignment of functional images across runs, tSNR, and manual segmentation quality. Even though the alignment between structural and functional data was suboptimal around the bottom of the frontal lobe area, it provides sufficient quality in about 95% of the cortex (Fig. 5A). Displacement errors in the residual parts of the brain were corrected by means of manual segmentation in the distorted T_1_-weighted EPI space (Fig. 5B). We found slight differences in B_1_-inhomogeneity patterns across sessions, possibly because of small variations in the positioning of the dielectric pads. Still, alignment across sessions was excellent without visually detectable residual motion (Fig. 5C). This alignment was especially important in the sense of averaging functional data across sessions. Our signal acquisition/preprocessing quality is summarized in tSNR efficiency (Fig. 5D, E). Based on common standards in layer-fMRI described (https://layerfmri.com/qa/), these efficiencies were “excellent”. Depicted values of tSNRs refer to single individual runs. Over 60 hours of manual correction on GM segmentation corrected crucial miss-segmentation (Fig. 5F, G). For example, the calcarine sulcus was partly missing, frontal areas were shifted due to mis-alignment, and deep layer groups were underestimated overall.

**Fig 5:**
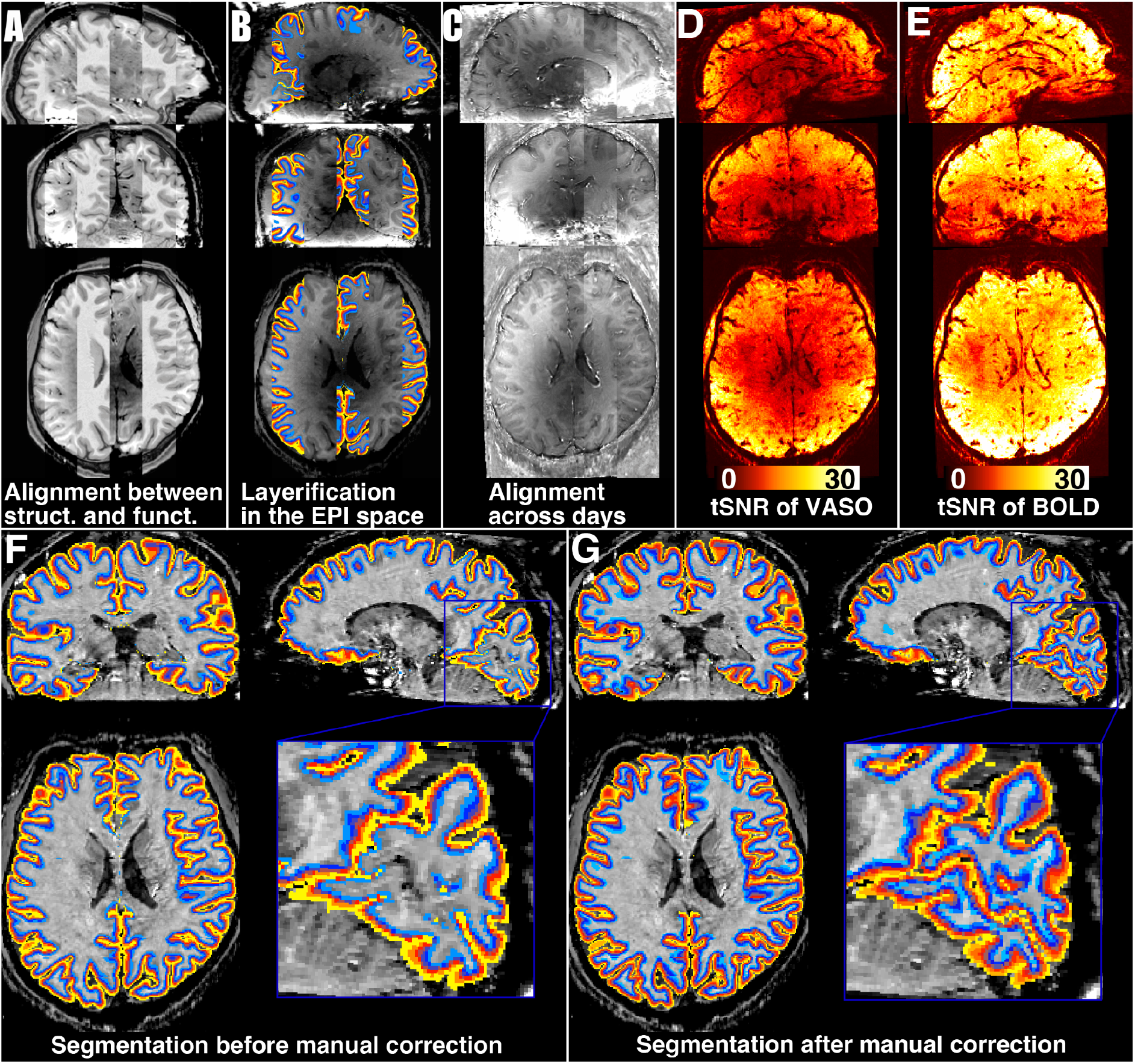
Preprocessing quality. For an animated version of this figure, please see here. Panel A: Non-linear alignment quality of a 3T MPRAGE to VASO EPI. Most of the brain is of usable quality. Panel B: Layerification quality in the EPI space. Panel C: Session-to-session alignment quality. The animated version of this figure cycles across individual sessions of multiple days. Note that the spatially heterogeneous brightness level of the T_1_-weighted EPI is a feature of variable flip angles in the MAGEC-VASO approach with spatially varying dielectric pads and does not affect the functional contrast evolution within each run. Panel D, E: tSNR efficiency of single run data in VASO (Panel D) and in BOLD (Panel E). BOLD time series show higher tSNR values than VASO. Average tSNR values were over 20 in most brain areas, which is excellent for a measure of sub-millimeter fMRI data (https://layerfmri.com/qa/). Panel F, G: Layerification before (Panel F) and after (Panel G) manual correction. Initially, parts of the calcarine sulcus were not classified correctly or only a few hundred micrometers thin. Frontal brain areas were shifted due to residual mis-alignment of the structural-functional image registration, and the layerification did not reach the deep layers everywhere. All these major imperfections were corrected by manual corrections (over 60h manual labor).

Figs. 6 to 11 exemplify the neuroscientific applicability of the provided data for a collection of popular fMRI analysis approaches. As such, Fig. 6 shows that the developed imaging methods are capable of capturing layer-dependent differences of common ICA networks. The Parieto-Frontal network, the Visual network, and the Auditory network show consistent topographical distributions across depths. On the other hand, the ‘default mode network’ is interestingly showing considerable deviations along the lateral direction of the cortex. We find that the visual and auditory networks show stronger z-scores in superficial and deep layers, compared to middle layers (Fig. 6). This finding might be related to the fact that neurons even in those uni-modal areas have more feedback synapses from cortical areas compared to feedforward input. It might also be associated with the fact that the participants could anticipate visual and auditory stimuli after seeing them many times already. When analyzing each session’s data separately, we see consistent layer signatures, demonstrating high repeatability (Fig. 7).

**Fig 6:**
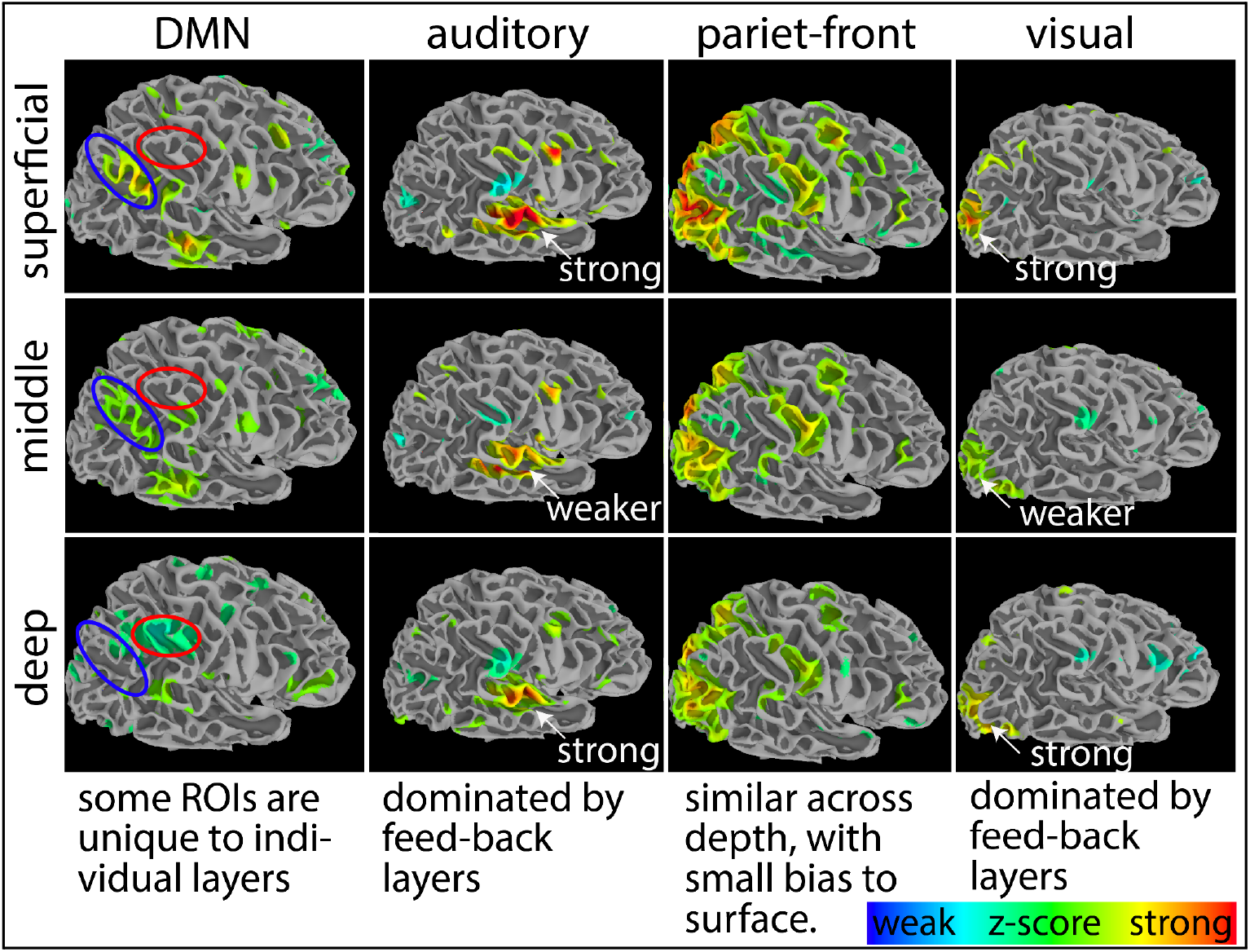
Layer-dependent differentiation of ICA-determined connectivity networks The purpose of this figure is to exemplify the neuroscientific usability of the provided data, beyond tSNR maps and strong tapping/checkerboard tasks. The surface projections of ICs for three layers. Four representative networks are selected. It can be seen that the Kenshu-dataset has sufficient functional sensitivity to extract common previously described brain networks without being limited by salt-pepper thermal noise. Here at 0.8mm resolutions. We find that the primary areas (e.g., visual and auditory cortices) are showing stronger z-scores for superficial and deep layers than for middle layers. The default mode network (DMN) stands out in the way that it seems to subdivide into different sub-ROIs for different layers. Activation in other networks (e.g., parietal-frontal) is relatively consistent across depths. The data shown here refer to VASO during movie watching (vein-contaminated BOLD data are not shown).

**Fig 7:**
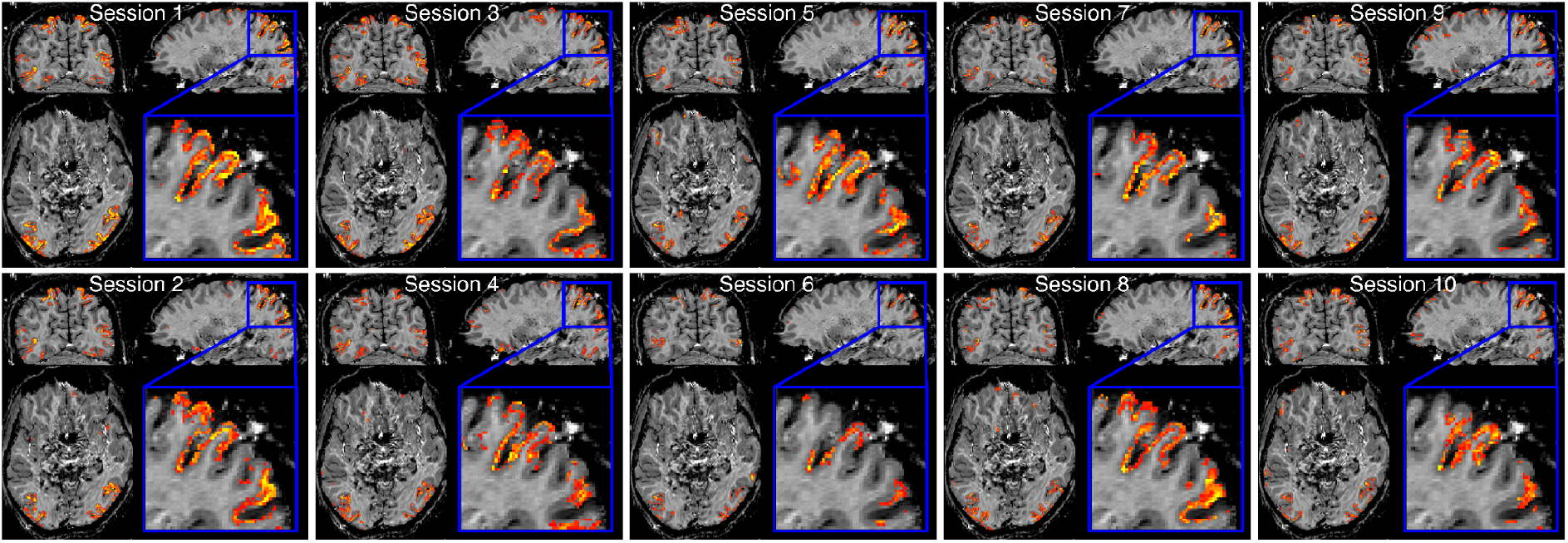
Replicability of movie networks across sessions. The ICA analysis of movie-watching data (as shown in Fig 6) provided signal traces of expected activation time courses for each network. Here we used the signal trace of the fronto-parietal network (representative for any other network) and used it as a regressor in a conventional GLM analysis. Each panel refers to a session average of 5 runs each. We can see consistent layer signatures for each independent day’s result. This supports the feasibility of whole-brain layer-fMRI to extract neural information in associative brain areas. The session-to-session repeatability is visible despite the fact that the participant did not have to follow a fixation task during free movie watching. The data shown here refer to VASO during movie watching (vein-contaminated BOLD data are not shown).

**Fig 8:**
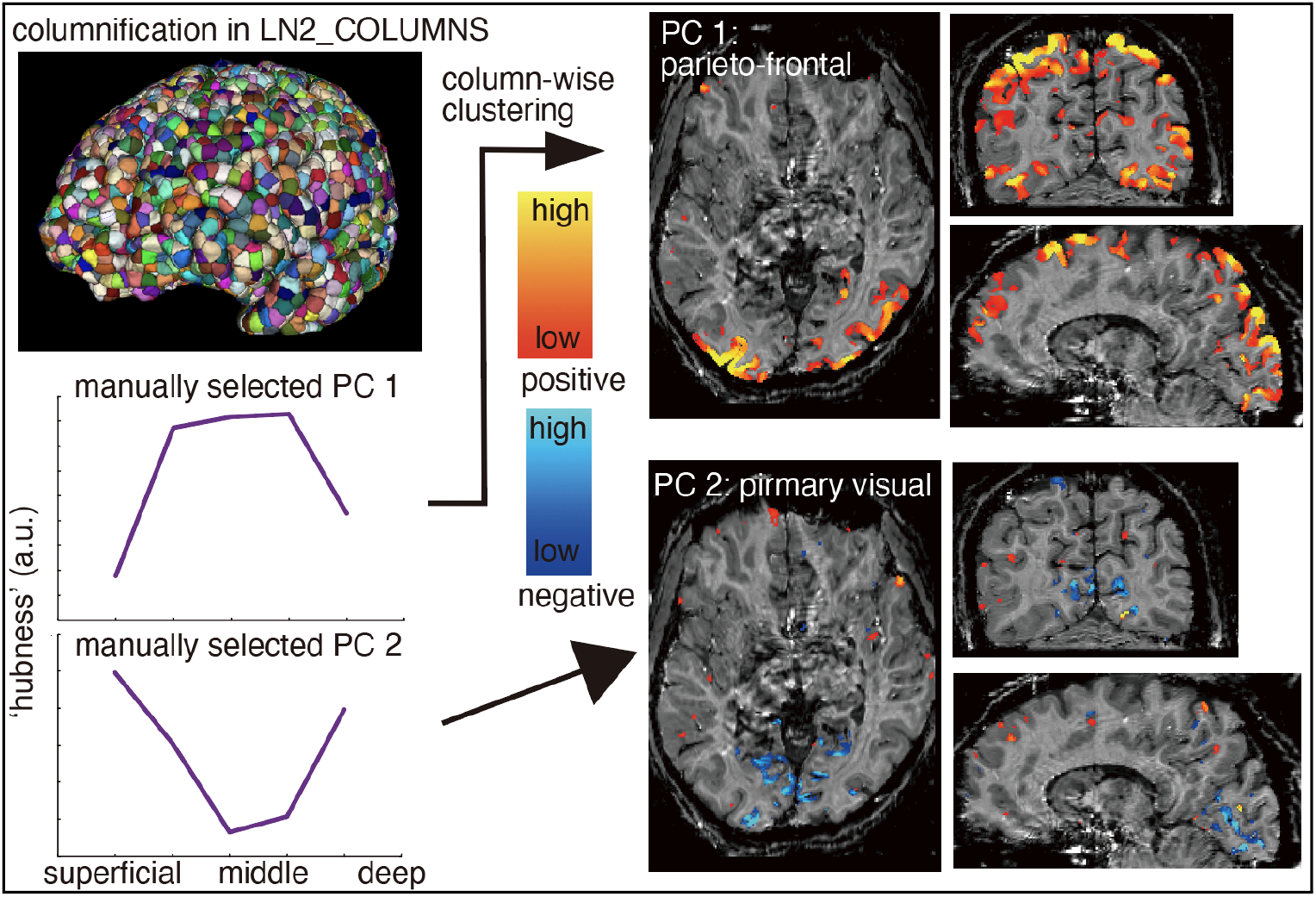
Network detection replicability and layer-profile analysis In this analysis, we first estimated the voxel-wise ‘hubness’ (AFNI’s 3dTcorrMap) of session-averaged time series. This measure indicates how much each voxel’s time course represents the overall fluctuations. Then, we divided the brain into 10,000 columnar ROIs with LayNii’s LN2_COLUMNS (top left). In some columns, hubness values were largest in middle layers and in some other columns hubness values were larger in superficial and deeper layers. We performed a PCA to group columns with similar layer-profiles in a data-driven way. This step does not impose hypotheses about expected profile shapes. The outcome of two principal components are depicted at the bottom left and exhibit feedforward-like (middle layer dominance) and feedback-like (superficial and deeper layer dominance) signatures. The right panel depicts the corresponding spatial maps of these profiles, respectively. It can be seen that they are confined to hemispherically symmetric brain networks. Hence, we think that these results exemplify the potential usability of whole-brain layer-fMRI protocols for neuroscientific research questions about laminar connectivity.

**Fig 9:**
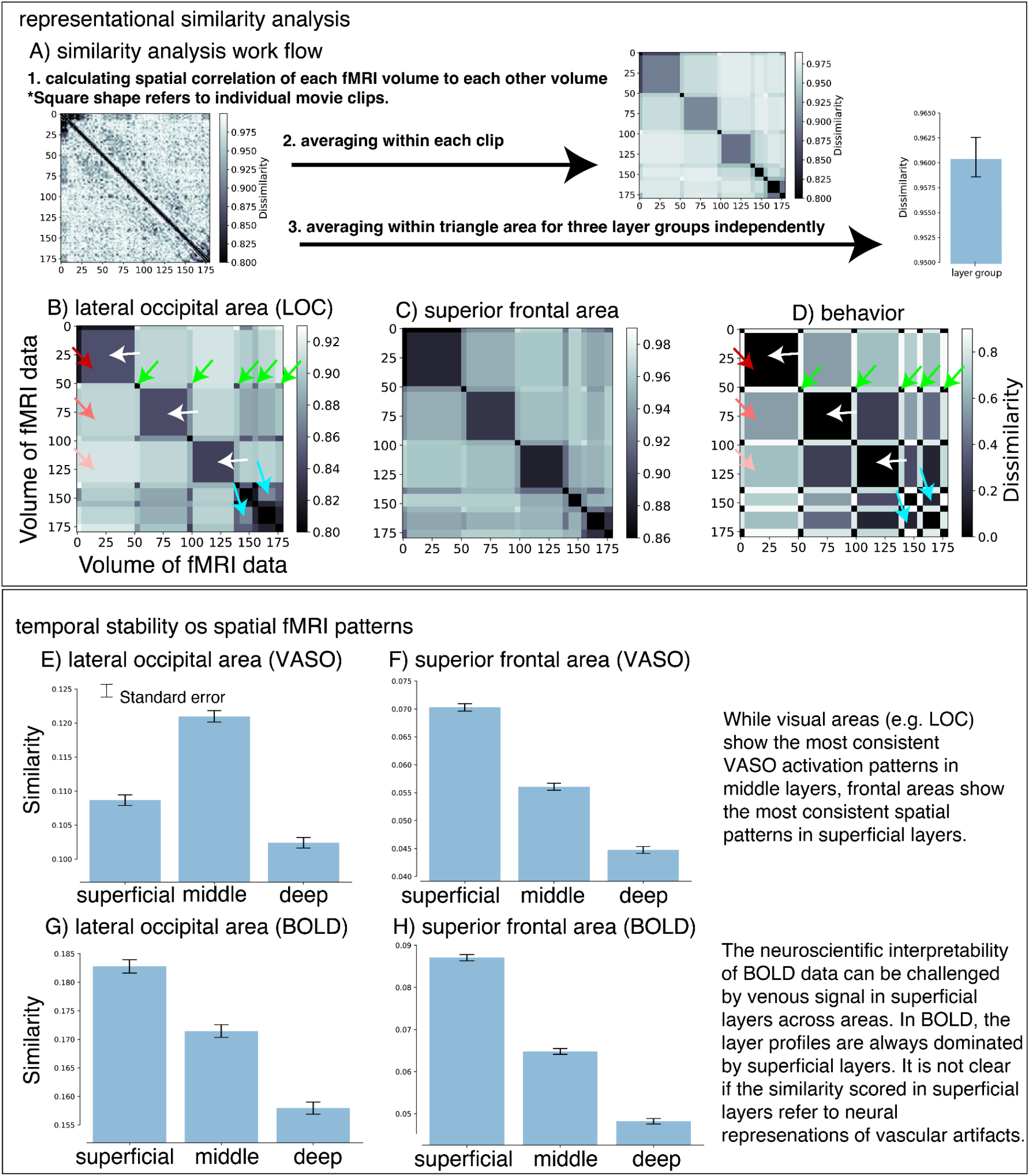
RSA and movie episodes (incl. rest) for fMRI and behavior Panel A schematically depicts the analysis workflow employed here. We calculated the RDMs by comparing the spatial similarity of each movie frame with each other movie frame in the time series. The representational similarity values (one minus Pearson’s r) are collected in a matrix form, which already shows indications of movie events (horizontal and vertical stripe features). For clarity, we averaged similarity values for each clip duration. Moreover, to see the layer profile or similarity consistency across time series, we calculated the mean and standard errors in each layer. Panel B-D depict these RDMs for representative brain areas of the lateral occipital area (panel B) and superior frontal area (panel C) (all other brain areas can be browsed here). As a reference, the behavioral similarity of each clip is also mapped in the same design (panel D). We find that the similarity structure of the brain data and the behavior matches for some ROIs. E.g. matrix features are similar in LOC and behavior (colored arrows). Such clear correspondence is not visible with other brain areas (e.g. frontal areas). Panel E-H: We also calculated the layer-specific similarity within brain areas. Here similarity refers to the overall brightness of the matrix in panels A-C. This metric does not reflect the matrix pattern across different movie clips but instead, it is a measure of the overall stability of the fMRI signal fluctuations across the entire run. We find that different ‘layer profiles’ are observed across different ROIs. It can be seen that VASO data exhibit different layer-profiles across ROIs, whereas BOLD seems to be always dominated from superficial layers. This might be due to large draining vein effects in BOLD. All panels combined suggest that whole-brain layer-fMRI data can provide plausible representational similarity results. We believe that this suggests that the whole-brain layer-fMRI protocols developed here can be a useful applicable tool for neuroscience research questions.

**Fig 10:**
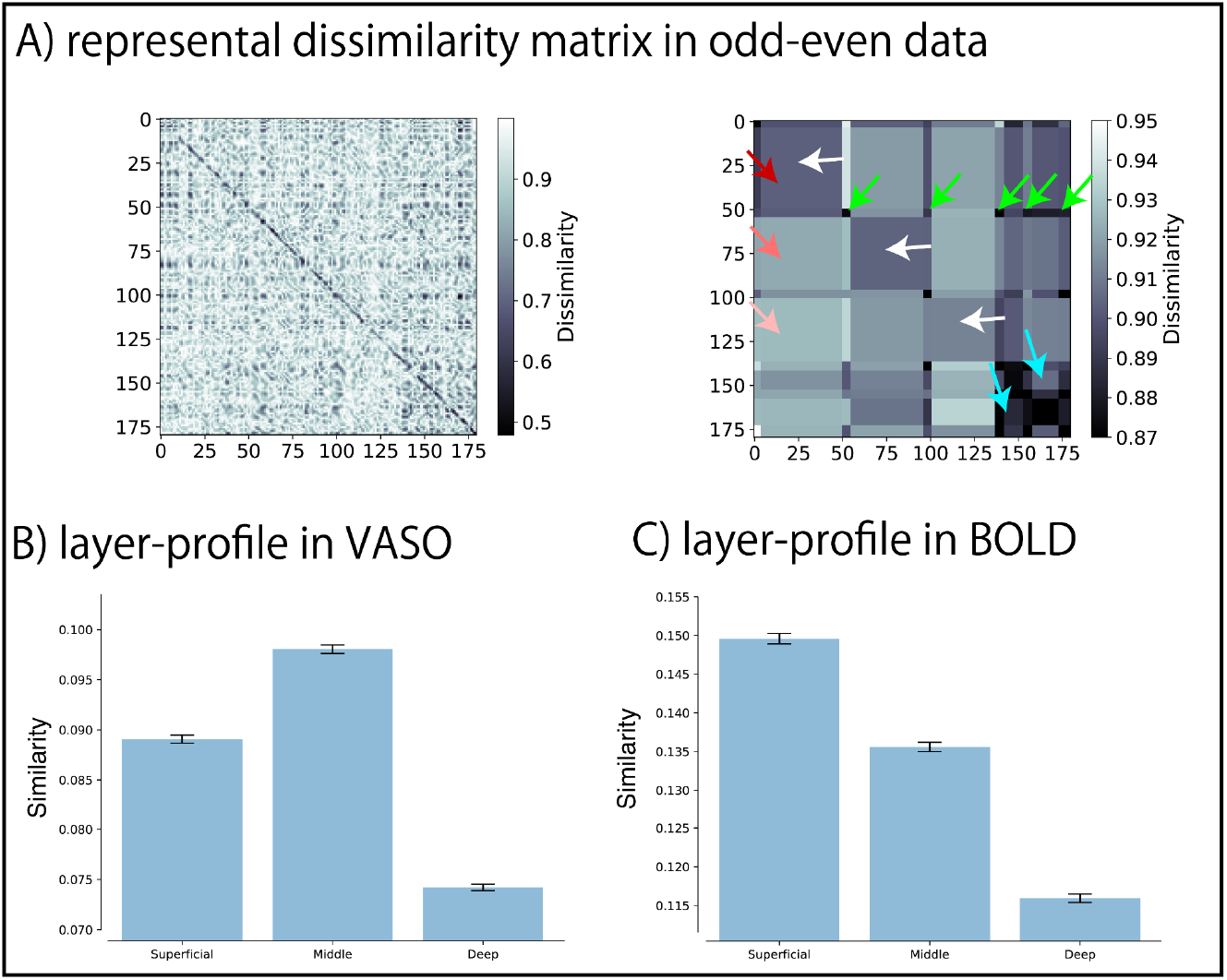
Replicating the RDM results of Fig. 9 by pooling data from independent sessions. RDMs were also estimated by comparing spatial similarities of independent data (even/odd sessions). It can be seen that the dissimilarity structure from Fig. 9 is replicated. Arrows point to replicable features. Namely, white arrows show that the similarity of spatial brain activation patterns for any given time frame and movie clip is similar across sessions. We interpret this result as a strong indication that the features in the similarity matrices are not dominated by volume-specific and run-specific MR noise. The low dissimilarity across rest periods (green arrows), the relatively reduced spatial dissimilarity across the final two fast-paced movie clips (blue arrows), and the successively increasing dissimilarity across the first three full movie clips (red arrows) is consistent with results in Fig. 9. Here, we present results in LOC, representative for all other brain areas. The same results for all other areas can be found here.

**Fig 11:**
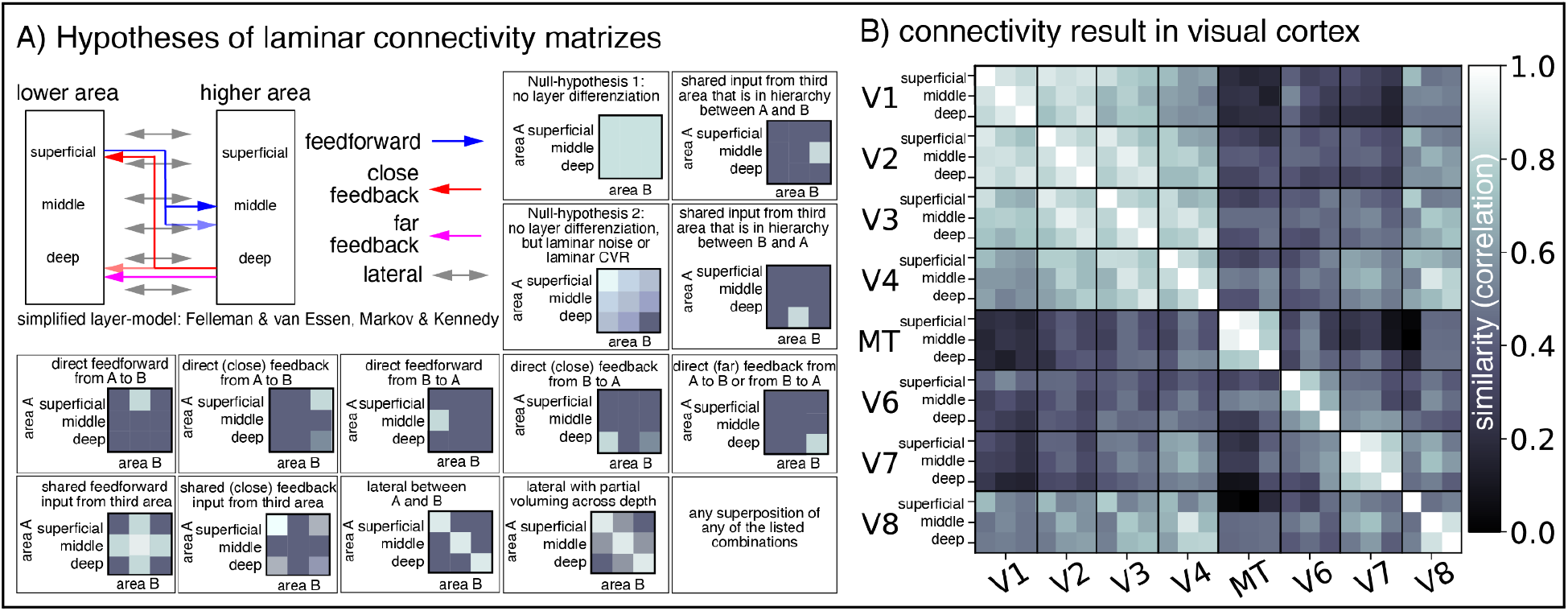
Functional connectivity (temporal similarity) analysis across layers and visual areas Panel A depicts hypothesized connectivity matrices. Based on the canonical microcircuitry model (Felleman and Van Essen 1991; Markov and Kennedy 2013), a collection layer-dependent connectivity matrizes could be expected. A subset of selected connectivity fingerprints is shown. Panel B depicts the empirical temporal similarity (connectivity) across three layer groups within and across visual areas V1-V8. Overall, we find high correlation within early visual areas V1-V4 compared to correlations among higher visual areas. Generally, the functional connectivity values vary more across areas than across layers. Focusing on the laminar fingerprints of ROI-pair sub-matrices, we find that correlations between superficial layers tend to be higher than other layers in most of the ROI-pairs. This might suggest shared feedback input. Aside from the diagonal submatrices (within layer connectivity) we find that almost all ROI pairs show laminar connectivity fingerprints that are different from the Null-hypotheses 1&2. This suggests that the layer-fMRI acquisition and analysis tools developed here can be useful to address questions of hierarchical neural connectivity across brain wide cortical networks. The Null-hypotheses refer to potential connectivity matrices that cannot be interpreted with respect to layer-specific differentiation of functional connectivity. The examples are: 1) all layers exhibit the same correlation values. 2.) correlation values are scaled versions of each other across cortical depths. E.g., due to depth-dependent variations of SNR, or due to depth-dependent variations of vascular reactivity.

Furthermore, we used column-wise PCA analyses of all layer profiles to explore data-driven approaches of consistent layer profiles across areas (Fig. 8). Doing so for movie-watching tasks, we find that columns that exhibit layer-profiles of inverted U-shapes (feedforward-like) tend to be stronger represented in the parieto-frontal area. On the other hand, for the experimental setup used here, we find that columns with U-shaped (feedback-like) layer-profiles tend to be strongly represented in visual areas. Seeing such clear distinctions of brain areas based on their layer profile shapes supports the value of laminar fMRI tools to inform neuroscientific interpretation of brain data. Based on the canonical microcircuit model (Felleman & van Essen 1991) each and every brain area (along the visual processing stream) is expected to contain both, feedforward and feedback inputs, respectively. However, for a given experimental task-condition is not clear, which of those pathways dominates. As such, it has been shown that in V1, middle layer (feedforward) input often dominates the overall ongoing neural activity for low-level visual stimuli (Self et al. 2019). Here we find that for repeated movie watching tasks and for measures of hubness, V1 columns are dominated from feedback layers.

We have also explored the usability of whole-brain layer-fMRI protocols for multi-voxel pattern analyses (MVPA). Specifically, we focused on representational similarity analyses. An example of an RDM for a representative brain area is shown in Fig. 9A. Indications of vertical and horizontal stripe-structures refer to events across movie clips. These movie event signatures are visible across the majority of brain areas and layers. Moreover, when calculating the mean within each clip, we see the movie structure more clearly. Specifically, we find that the spatial representational pattern is more similar within movie clips as opposed to across-movie clips (diagonals are darker in Fig 9A-C, white arrows). Results in Fig. 9 refer to representative brain areas of lateral occipital cortex (LOC) and superior frontal cortex. Here, we decided to include LOC as an example of visual processing that is expected to be highly coupled to the movie timing, while having rather nonspecific receptive field sizes, effectively mitigating dependencies of visual gaze. We also included an example association area as reference (superior frontal cortex). Results of all other brain parcels that are part of the Glasser atlas, are available here (https://zenodo.org/record/6882801).

The visualization of layer-fMRI brain data in the form of RDMs allows comparison with behavior. As such, it looks like that the matrix of subjectively perceived movie-clip-similarities matches RDMs of LOC quite well (see colored arrows). As such, the last two movie clips are the most similar ones (blue arrows). Those are fast-paced repeat clips. Also, the first three movie clips are successively increasing their dissimilarity (red arrows). And lastly, movie clips are more similar to movie clips than rest periods (green arrows). This correspondence is brain-area-specific and not as clearly visible in frontal areas. As RSA, calculating correlations between behavior data and VASO/BOLD data, the correlations were significant (Bonferroni-corrected) in most of the ROI (30 out of 31 in VASO and 29 out of 31 in BOLD). Moreover, we found that VASO data have higher correlations than BOLD data in most of the ROI with significant correlations between behavior data and VASO/BOLD data (28 out of 29) (E.g. VASO: 0.106, BOLD: 0.046 in LOC all layers).

We have also calculated the similarity values across layers. We find that the temporal stability of the spatial pattern in LOC is highest in the middle layers, while most frontal areas showed the highest similarity values in superficial layers.

Note that the results in Fig. 9 refer to spatial similarity patterns between all combinations of movie frames for volume time series that had been averaged across all 51 runs. This means that the diagonals of the RDM’s represent the spatial correlation of the fMRI brain data with itself. Thus, the diagonal similarity values might be overestimated by spatial patterns of random noise. Furthermore, this way of comparing within-run similarities might also include potential global biases of run-specific non-linear scanner drifts and other sources of run-specific temporal autocorrelation. This means that run-specific imaging artifact modulations across time could globally inflate similarity values. Since layer-fMRI acquisition protocols are quite demanding on the scanner hardware, such temporal artifacts and hardware instabilities are somewhat expected for layer-fMRI. In order to investigate if and how much the similarity patterns in the RDMs can be affected by such potential biases, we replicated the analysis with a different data pooling strategy. Namely, we pooled the runs into two separate datasets of odd and even sessions (see method section for more details) and then repeated all the following steps as described above. We find that similar matrix features were shown, independent of the data pooling strategy (compare arrows in Fig. 9 and 10). This suggests that the updated data pooling is not a result of correlating spatial structure of each time point with itself (including structured noise and neural representations).

Taken together, the results in Figs. 9-10 suggest that the layer-fMRI protocols developed in this study can facilitate such data-driven volume-by-volume representation similarity analyses.

We also explored analysis strategies that are based on the temporal similarity of fMRI time courses (a.k.a. functional connectivity). Due to the neuroscientifically established layer-dependent structural connectivity in the early visual cortex (Felleman and Van Essen 1991), we used visual areas V1-V8 as a ‘testbed’ to explore the usability of the developed protocols (Felleman and Van Essen 1991; Markov and Kennedy 2013; Jia et al. 2020; Yang et al. 2021). Functional connectivity matrices and potential connectivity hypotheses are depicted in Fig. 11. We find indications of various laminar feedforward and feedback signatures (Fig. 11 Panel A). There are indications of feedback signatures in: V1-V2, V1-V3, V1-V4, and V1-MT. Also overall, we find the largest similarities between superficial layers compared to deeper layers (Guidi et al. 2020; Huber et al. 2017; Deshpande, Wang, and Robinson 2022; Pais-Roldán et al. 2020). This might indicate that feedback signals dominate the temporal signal fluctuations during repeated movie watching. The results presented in Fig. 11 suggest that the layer-fMRI protocols developed here are usable to capture layer-specific differentiates of functional connectivity. This implies that the tools developed here can be valuable to address research questions of directional information flow across the entire cortex.

In recent years, the research field of layer-fMRI methods development has invested some efforts into laminar vascular deconvolution models (Markuerkiaga, Barth, and Norris 2016; Akbari et al. 2021; Havlicek et al. 2015; Heinzle et al. 2016; Huber et al. 2021). The purpose of these models is to account for unwanted venous biases of layer-fMRI signals in conventional gradient echo BOLD. Until now, these models are implemented and validated for the specific case of the primary visual cortex. It is still not clear, however, how translatable their application is to other areas with potentially different baseline vascular physiology. For example, venous baseline blood volume is believed to increase towards the cortical surface. And small changes in the slope of assumed venous baseline blood volume increases can result in substantially different results of ultimate deconvolved laminar activity changes (Huber et al. 2017). Thus, for future applications of laminar deconvolution models beyond the early visual cortex, they need to be extended to a wider range of ROI-specific differences in venous blood volume. We believe that the whole brain dataset presented in this work can be useful to investigate areal differences of these laminar angio-architectonic characteristics. The dataset contains both BOLD and VASO data across all brain areas. Thus it can serve as an atlas-like reference to fine-tune model parameters of laminar deconvolution analyses for any cortical area of interest. Here we estimated measures of venous baseline blood volume distributions by means of previously established and calibrated VasA methods (Kazan et al. 2017). Fig. 12 depicts three projections of the 3D atlas and layer-profiles of representative brain areas (LOC and SFC). It can be seen that pial voxels have larger values of cerebral vascular reactivity compared to deeper GM voxels. We find this increase in reactivity towards the surface in ≈170 of all 180 cortical areas out of the Glasser atlas. Most brain areas show layer-profiles that increase supra-linearly towards the surface. The few exceptions that do not show a clear increase are referring to ventral limbic areas.

**Fig 12:**
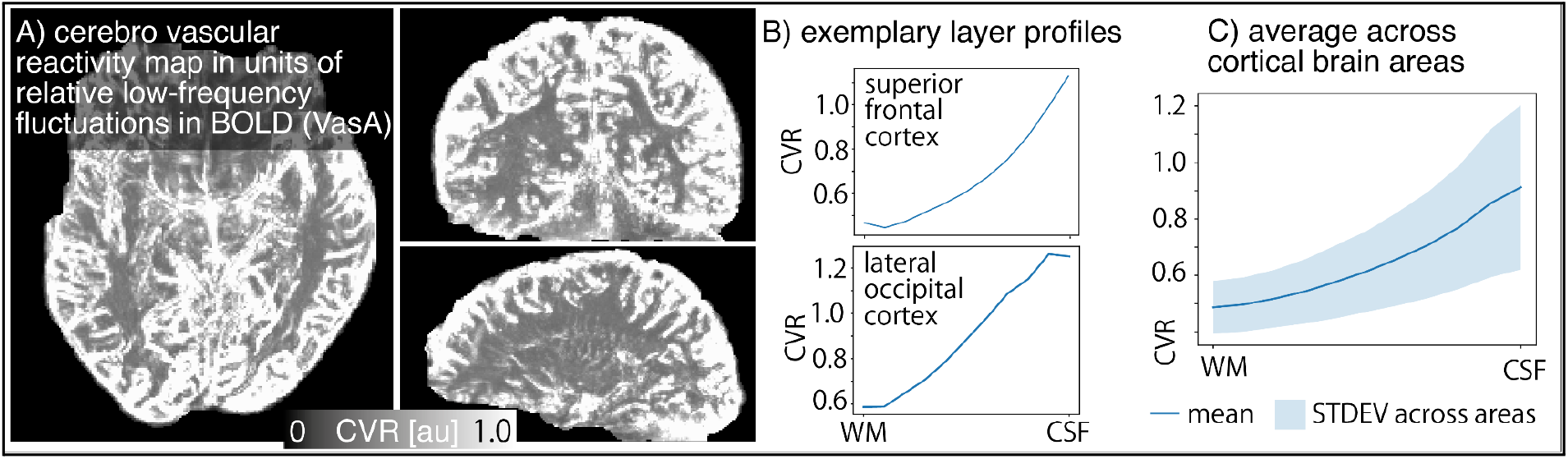
Atlas of laminar vascular reactivity estimates Panel A depicts representative projections of vascular reactivity estimates at 0.8mm resolution. It can be seen that high CVR values are following the cortical anatomy and possibly macro-vascular structures. The full 3D dataset is available on Openneuro. Panel B depicts representative layer profiles of CVR values across cortical depth. For consistency with previous figures, exemplary areas are superior frontal and lateral occipital cortex. It can be seen that CVR estimates in superficial layers and pial voxel are about twice as high as deeper layers. This is consistent with almost all other cortical brain areas. 2D-plots and tables of all layer profiles of all brain areas in the Glasser atlas are available on Openneuro (https://openneuro.org/datasets/ds003216). Panel C depicts the average and standard deviations (STDEV) across brain areas. It can be seen that the supra-linear increasing CVR values towards the surface are relatively consistent across brain areas.

## Discussion

Previously described layer-fMRI acquisition protocols with limited coverage can address questions of directional information flow during cognitive mental processes (Finn et al. 2019; Schneider et al. 2019) in health and disease (Stephan et al. 2019; McColgan et al. 2020; Haarsma, Kok, and Browning 2020). However, small imaging coverages are not straightforwardly standardizable and do not allow layer-fMRI to fulfill its promise to capture functional directional connectivity. Namely, in order to capture neural information flow during task and rest, multiple brain areas need to be captured at the same time; i.e., the brain area(s) that send neural information and the brain area(s) that receive the information need to be sampled concomitantly. For this, larger coverages are necessary. This study on the development of whole-brain layer-fMRI acquisition and analysis protocols provides the tools that allow the field to overcome these limitations.

We describe an openly accessible, vein-free, whole-brain layer-fMRI dataset for developing and benchmarking functional laminar analysis methodologies. Along with the dataset, we share layer-fMRI-optimized and highly vetted acquisition and preprocessing analysis protocols that can be openly used by the neuroimaging community.

### Brain wide functional connectivity in layer-fMRI: investigating temporal similarity of fMRI signal changes across areas

Metrics of functional connectivity with fMRI have the potential to be informative of feedforward or feedback dominated neural information flow. Early attempts of layer-fMRI connectivity studies have been somewhat limited to relatively small field of views, constrained to individual brain systems (Polimeni et al. 2010; Guidi et al. 2020; Huber et al. 2017; Huber, Finn, et al. 2020). More recent advancements in data sampling approaches, MR-contrast generation strategies, and confidence of the laminar signal interpretability allowed proof-of-principle extensions of layer-fMRI connectivity to larger FOV (Sharoh et al. 2019; Huber et al. 2021; Deshpande, Wang, et al. 2022; Pais-Roldán et al. 2020; Yun et al. 2022). In this work, we aim to help the layer-fMRI community in building tools to make such whole-brain layer-fMRI connectivity protocols usable for neuroscience application studies. Until now, this is limited by multiple aspects:

- There are no established preprocessing analysis tools for whole-brain layer-fMRI connectivity data. The extremely high accuracy requirements of layer-fMRI, combined with the challenging artifact level and non-linear geometric distortions does not allow straightforward applications of conventional preprocessing pipelines. Here, we described how to mitigate this challenge in two ways. 1.) We describe our attempts of building a complete analysis pipeline that fulfills layer-fMRI quality standards (Figs. 1-5). 2.) We share a benchmarking dataset which can be used as a ‘testbed’ for developing and benchmarking new analysis tools (Figs. 8-10).
- The test-retest reliability of layer-fMRI’s detection sensitivity to capture large-scale connectivity networks is still unclear. In Fig. 7, we characterize and repeatability of such whole-brain connectivity data up to 10 sessions.
- We exemplify the neuroscientific applicability of the whole-brain layer-fMRI connectivity protocols developed here. Specifically, in the results presented in Figs. 6-8, 11, we confirmed our neuroscientific applicability of common connectivity analysis tools, including: ICA, hubness mapping, and temporal correlation matrices.

### Representational similarity in whole-brain layer-fMRI: investigating spatial similarity of fMRI signal changes across TRs

In order to investigate the neuroscientific usability beyond temporal similarities (a.k.a. resting-state connectivity), we explored the usability of RSAs. Based on previous work on repeated movie watching at lower resolutions (Baldassano et al. 2017), we expected to see temporal features of movie events across representational similarity matrices. E.g., we expected to obtain different spatial similarity scores between rest clips and movie clips, and we expected to see different spatial similarity scores between pairs of movie clips. This would indicate which movie clips are more similar to each other compared to other movie clips. Exploring laminar differentiations of representational similarity strengths can then be helpful to build hypotheses about the underlying neural feedforward and feedback mechanisms that are involved in these representations. If such temporal features in the RDMs are not visible, this suggests that layer-fMRI is too SNR limited for such high-level analysis frameworks.

Figures 9-10 show examples of the temporal event structures within RDMs, especially for movie clip-averaged data. This indicates that the layer-fMRI protocols developed here are indeed applicable for RSA. In VASO data, we found that in different brain areas the representational similarity strength is differentially dominated by feedforward and feedback layers, respectively. E.g., while the visual area LOC shows feedforward signatures, superior frontal areas have highest representational similarity strengths in cortico-cortical feedback input layers. In BOLD data however, we did not see such clear differentiation across areas. Instead, almost all brain areas are dominated by largest representational similarity strengths in the superficial layers, which might be caused by its sensitivity of large draining veins.

Until today, such RSAs have not been applied on sub-millimeter layer-fMRI data. Thus, it is not yet established if they might be biased by run-specific, temporal signal artifacts. In order to confirm that the spatial similarity results are interpretable as movie synchronized neural representations, we validated the results with independent pairs of datasets (Fig. 10). We find that the temporal event structures in the RDMs and the corresponding layer-profiles are qualitatively identical. This supports the neuroscientific usability of the protocols and data described here.

### Temporal sampling efficiency of whole-brain layer-fMRI?

The whole-brain layer-fMRI acquisition protocol developed and used here has a volume acquisition time of 5.1s. While such volume TRs are longer than typical fMRI acquisition protocols, the data acquisition rate is considerably higher than conventional layer-fMRI protocols. Namely, the EPI protocol of this study samples more than 1,000,000 voxels per second. While faster acquisition protocols with volume TRs down to 2.3s are possible (Fig. 3), the correspondingly higher noise level results in overall lower tSNR efficiency.

Here we chose to use this relatively long volume TR based on considerations to maximize functional detection sensitivity. Namely, the power-spectrum of functional connectivity fluctuations as used here have their its highest contrast-to-noise-ratio around 0.01Hz - 0.1Hz (Biswal et al. 1995; Lowe, Mock, and Sorenson 1998; Cordes et al. 2000; Fransson and Marrelec 2008; De Luca et al. 2006; Wu et al. 2008). Thus, being in the thermal noise limited regime of sub-millimeter fMRI, it is advised to optimize the acquisition protocol for this frequency regime too. Furthermore, there is emerging evidence that for connectivity-based analyses with resting-state or movie-watching data, fast sampling is not vital. As such:

- (Airan et al. 2016) showed that the number of data points is less critical than acquisition time. Specifically implying that short TRs cannot make up for short scan durations (specifically Fig. 4, top panel).
- (Huotari et al. 2019) showed that brain networks can be sampled across TRs across several orders of magnitude without losing the ability to capture conventional brain networks.
- Jason Druzgal et al. had shown that connectivity-based fingerprinting of whole-brain movie-watching paradigms are relatively constant across TRs. Only for TRs longer than 10s, the fingerprinting accuracy starts to decline.
- (Birn et al. 2013) have also shown that the scan duration is more important than sampling rate when holding the number of data points constant (see especially Fig. 4, with TRs up to 5.2s).

While meaningful neural and corresponding vascular activation modulations are happening across a wide range of temporal frequencies (Godlove et al. 2014; Sotero et al. 2015; Ma et al. 2016; Lewis et al. 2016), connectivity-based fMRI fluctuations follow the pattern of ‘scale free dynamics’ (He 2011). This means that focusing on one specific frequency window is expected to provide results that are largely representative of functional connections at any other temporal scale. Ongoing work in combining the whole-brain layer-fMRI approaches with further advanced hardware (Feinberg et al. 2022; Beckett et al. 2022) and MR-reconstruction (Yun et al. 2022) will become important to confirm this temporal invariance.

### Future methodological developments

Previous large coverage layer-fMRI acquisition protocols exist and exemplify that current acquisition tools are -in principle-sensitive to capture depth-dependent connectivity differences (Finn et al. 2021; Bandettini et al. 2021; Huber et al. 2021; Yun et al. 2022; Pais-Roldán et al. 2020; Sharoh et al. 2019; Chang et al. 2022; Deshpande, Zhao, et al. 2022; Müller et al. 2021). Here, we move beyond the proof-of-principle concept and characterize the stability of such acquisition protocols, we developed a preprocessing protocol, and we explored the neuroscientific usability of whole-brain layer-fMRI. In this work, we could show that whole-brain layer-fMRI provides repeatable results across many sessions and that it can be combined with less controlled, naturalistic tasks. While this opens a large range of potential application cases, further methodological acquisition and preprocessing developments will be helpful still.

- **NORDIC-PCA**: Due to the high spatial resolution and relatively liberal GRAPPA acceleration used here, the raw data are in the thermal noise limited regime. This means that the application of NORDIC-PCA denoising (Vizioli et al. 2021), might be helpful to reduce the number of necessary runs and/or might allow even further acceleration. Each run of the Kenshu dataset contains two noise volumes that are acquired and shared for the purpose of potential future NORDIC-PCA. So far, we did not apply NORDIC denoising, because it is not yet established for tasks beyond on-off (e.g. blocked) stimulation periods outside of exploratory use. The Gaussian-like temporal activity modulations of naturalistic tasks of the Kenshu dataset might require specific parameter optimization in NORDIC in order to ensure that no activation of interest is lost.
- **Extension across participants:** While the study described here refers to approx. 60 hours worth of data (including all pilot sessions), the main 51-run data are collected from one single participant. We deliberately focused all scanning funds on one participant in order to capture the test-retest reproducibility of the methods (not across people). Thus, this dataset cannot be used to parameterize other sources of variance that might arise from individual differences. Future work is necessary to extend the dataset to more participants.
- **Extension to newer scanner platforms:** The Kenshu dataset was acquired on a ‘classical’ SIEMENS MAGNETOM 7T scanner. The corresponding scanner console is still running on Windows Xp and will be outdated in the coming years. The vendor that dominates the 7T MRI market predicts that 70% of the scanners will be upgraded to a newer ‘Terra’ console by the end of 2022. Future work is needed to translate the acquisition protocols developed here to whole-brain VASO sequences that are readily available across consoles (Stirnberg 2021; Huber 2021). Such sequence approaches introduce further flexibility of segmentation across inversion-recovery cycles.
- **Increasing spatiotemporal resolutions with advanced hardware:** The Kenshu dataset is acquired at volume TRs of 5.1s and spatial resolutions of 0.84mm. Compared to other layer-fMRI studies in the field, this is considered to be rather liberal. Higher spatiotemporal resolutions would be advantageous to better separate different layer groups at faster neuronally relevant time scales. Ongoing research on whole-brain layer-fMRI VASO with high RF channel counts and high-performance gradients is being conducted (Beckett et al. 2022; Feinberg et al. 2022).

### What this dataset can be useful for

We believe that the dataset presented here will serve the neuroimaging community in multiple aspects. Most importantly, this dataset allows developers of fMRI pre-processing and post-processing tools to adjust and extend their methodology to become applicable for layer-fMRI. Specifically,

- In ongoing studies, (Akin et al. 2022) are using the Kenshu dataset to explore the feasibility of extending layer-fMRI to capture cortico-subcortical connections.
- In ongoing studies, (Klein et al. 2022) take advantage of the Kenshu dataset to measure functional connectivity of large scale cortical brain areas. Namely, the functional connectivity between V4 and frontal areas.
- Furthermore, Tomas Knapen and his students are using the Kenshu dataset as a testbed to develop connectivity-derived retinotopy representations across brain areas and cortical layers, following their previous work at lower resolutions (Knapen 2021).
- The Kenshu dataset is also turning out to be useful as an exemplary dataset for showcasing the working principles layer-fMRI viewing tools in AFNI, such as the tools SurfLayers (Torrisi et al. 2021) or clipping planes for layer-fMRI (Lauren et al. 2022).
- The dataset used here was also helpful for Logan Dowdle to test the robustness of non-linear motion-correction and alignment protocols. Such work is specifically important in order to address issues of geometric distortions, which are particularly challenging in layer-fMRI protocols that are limited by low readout bandwidths.
- While layer-fMRI and VASO is still in the process of becoming an established standard neuroscience tool, at recent BrainHack events (Gau et al. 2021), Remi Gau has used the Kenshu dataset to exemplify how one could store layer-fMRI VASO data in an extended BIDS data storing standard. Link to this VASO fMRI BIDS demo: https://gin.g-node.org/RemiGau/ds003216/src/bids_demo.
- In ongoing sequence development projects, Yuhui Chai is taking advantage of the developed k-space acquisition procedures (acquisition protocols and sampling patterns) for whole-brain sub-millimeter EPI acquisition with MT-prepared and VAPER prepared contrasts (Chai et al. 2021).
- Preliminary accounts of this dataset have been used to implement, calibrate, and validate resolution biases in layerification algorithms (Figure 4 in Huber et al. 2021).
- Stability estimates derived from this dataset have been useful to discuss the challenges and opportunities of future mesoscopic brain imaging (Fig. 2 in Bandettini et al. 2021).

Moreover, we think that this work is particularly useful to facilitate the translation of MR-physics tools to the field of cognitive neuroscience. In the history of the layer-fMRI field, it has taken approx. two decades from the first proof-of-principle papers in 1999 (Menon and Goodyear 1999) until the number of application papers exceeded the number of methods-focused papers (www.layerfmri.com/papers). This was partly due to the fact that MR-physicists focused more on pushing the limits of their methods than working on the user friendliness of their workflows. In this work, we take advantage of being a multidisciplinary author team consisting of MR physicists and computational neuroscientists. We sought to translate high-end MR-methods towards neuroscientific application studies by applying a collection of trendy analyses that seem to be popular in current neuroscientists research (e.g., the RSA, functional connectivity analysis). We feel encouraged to learn how ongoing studies are using the Kenshu dataset for even further higher-order applications and we are looking forward to seeing what kind of new information about the brain can be captured with the developed protocols.

## Summary and Conclusion

In this work, we aimed to serve the field of layer-fMRI and help build tools for future routine whole-brain layer-fMRI in application-based neuroscience research. Previous proof-of-principle work has already shown that large-coverage layer-fMRI can capture laminar variations of functional connectivity. Here, we aimed to move one step further and work towards optimizing whole-brain layer-fMRI protocols to make them a neuroscience research tool with established characterizations of reliability and usability. We have developed publicly available sequence, acquisition, and processing pipelines for whole-brain layer-fMRI. And we are sharing an extensive multi-contrast whole-brain layer-fMRI dataset for the purpose of benchmarking future methods development: The Kenshu dataset. In this article, we describe the quality of the acquisition and preprocessing methodology in terms of tSNR efficiency and test-retest reliability across 10 sessions. Furthermore, we exemplify the usefulness of these protocols and data by applying a series of popular post-processing analyses as they are commonly used in cognitive neuroscience fMRI studies. This includes connectivity analyses, RSAs, GLM analyses, etc. We believe that this work paves the road for measuring depth-dependent directional functional connectivity across the entire brain in routine neuroscience application studies.

### Further material and Data/Software availability

- Video descriptions of various stages of this study are available.
  - 9 min video showing ‘hands-on’ that there is a layer-specific functional connectivity signal that differs across cortical depth: https://youtu.be/Dzde7ELcsNg (2020).
  - 12 min video showing that this signal is not stable enough for neuroscience practice: https://youtu.be/Zm8bTfk_IpQ (2021).
  - 5 min video describing the acquisition, the analysis, and the results of the main 50-run dataset: https://youtu.be/plF3K5VqXHU (2022).
- All raw and pre-processed data are stored in the Brain Imaging Data Structure (BIDS) (Gorgolewski et al. 2016) format and openly accessible on OpenNeuro (Markiewicz et al. 2021): https://openneuro.org/datasets/ds003216.
- This sequence is available at SIEMENS’ C2P ‘app store’ Teamplay https://teamplay.siemens.com/.
- An extensive application-focused description of the sequence and how to use it can be found in the users’ manual page: https://layerfmri.com/ss-si-vaso-sequence-manual.
- Full list of scan protocol parameters: https://layerfmri.page.link/WBprotocol.
- Scientific report on pilot experiments: https://doi.org/10.5281/zenodo.6860807
- Layerification analysis software developed in-house is available on Github (https://github.com/layerfMRI/LAYNII).
- Specific application scripts used here are available on Github: https://github.com/kenshukoiso/Whole_Brain_Project.

## Help with scanning

These data were acquired with the kind support of Scannexus at their 7T MAGNETOM ‘classic’ SIEMENS scanner. We thank Mathilde Kennis for being ‘on-call’ in case of emergencies during the scans that were all conducted after-hours between 5pm and 11pm. We thank Steen Moeller for advice regarding how to include the noise volumes at the end of the time series for the potential future usage of NORDIC-PCA denoising preprocessing. We thank Sri Kashyap for guidance with the dielectric pads. We thank Miriam Heynckes for guidance with the auditory delivery inside the scanner with the Sensimetric system. We thank Emily Ma for help with field monitoring with the Skope field camera.

## Funding

Kenshu Koiso is funded by the Japan Public-Private Partnership Student Study Abroad Program (TOBITATE! Young Ambassador Program), by Japan Student Services Organization (JASSO), and by UEC fund.

Laurentius Huber was funded by the NWO VENI project 016.Veni.198.032.

Benedikt Poser is partially funded by the NWO VIDI grant 16.Vidi.178.052, by the National Institute for Health grant R01MH/111444 (PI David Feinberg) and by the H2020 FET-Open AROMA grant agreement no. 88587.

Yoichi Miyawaki is funded by JSPS KAKENHI (20H00600, 18KK0311).

Rainer Goebel and Omer Faruk Gulban are financially supported by Brain Innovation.

### Ethics

The scanning procedures have been approved by the Ethics Review Committee for Psychology and Neuroscience (ERCPN) at Maastricht University, following the principles expressed in the Declaration of Helsinki.

### Task

We thank Emily Finn for guidance regarding the application of naturalistic tasks and helpful comments about the optimal sampling rate (TR). Movie stimulus data were provided by the Human Connectome Project, WU-Minn Consortium (Principal Investigators: David Van Essen and Kamil Ugurbil; 1U54MH091657) funded by the 16 NIH Institutes and Centers that support the NIH Blueprint for Neuroscience Research; and by the McDonnell Center for Systems Neuroscience at Washington University.

## Diversity Statement

The Kenshu dataset has been acquired at 7T. Such Ultra-high field scanners are limited to ≈100 privileged MRI centers around the world. By sharing this dataset publicly, we hope we will allow a wider community to benefit from this technology. In ongoing research, we are currently aiming to also extend the whole-brain layer-fMRI imaging methodology to more readily available 3T scanners (Huber et al. 2022).

Recent work in several fields of science has identified a bias in citation practices such that papers from women and other minorities are under-cited relative to the number of such papers in the field (Dworkin et al. 2020). In the human layer-fMRI community, the average gender citation bias is 84% male, 15% female (https://layerfmri.com/papers/). We obtained the gender of the first author of each reference. By this measure (and excluding self-citations to all authors of our current paper), our references contain 48 (74%) male first and 17 (26%) female first. We look forward to future work that could help us to better understand how to support equitable practices in science.

## References

Airan, Raag D., Joshua T. Vogelstein, Jay J. Pillai, Brian Caffo, James J. Pekar, and Haris I. Sair. 2016. “Factors Affecting Characterization and Localization of Interindividual Differences in Functional Connectivity Using MRI: Individual Differences in Functional Connectivity.” Human Brain Mapping 37(5):1986–97. doi: 10.1002/hbm.23150.

Akbari, Atena, Saskia Bollmann, Tonima S. Ali, and Markus Barth. 2021. Modelling the Depth-Dependent VASO and BOLD Responses in Human Primary Visual Cortex. preprint. Neuroscience. doi: 10.1101/2021.05.07.443052.

Akin, Burak, Kenshu Koiso, Richard Klein, Daniel A. Handwerker, Laurentius (Renzo) Huber, and Peter A. Bandettini. 2022. “Laminar Level Subcortico-Cortical Interactions during Naturalistic Movie Viewing.”

Allen, Emily J., Ghislain St-Yves, Yihan Wu, Jesse L. Breedlove, Jacob S. Prince, Logan T. Dowdle, Matthias Nau, Brad Caron, Franco Pestilli, Ian Charest, J. Benjamin Hutchinson, Thomas Naselaris, and Kendrick Kay. 2022. “A Massive 7T FMRI Dataset to Bridge Cognitive Neuroscience and Artificial Intelligence.” Nature Neuroscience 25(1):116–26. doi: 10.1038/s41593-021-00962-x.

Avants, Brian B., Nicholas J. Tustison, Michael Stauffer, Gang Song, Baohua Wu, and James C. Gee. 2014. “The Insight ToolKit Image Registration Framework.” Frontiers in Neuroinformatics 8. doi: 10.3389/fninf.2014.00044.

Baldassano, Christopher, Janice Chen, Asieh Zadbood, Jonathan W. Pillow, Uri Hasson, and Kenneth A. Norman. 2017. “Discovering Event Structure in Continuous Narrative Perception and Memory.” Neuron 95(3):709-721.e5. doi: 10.1016/j.neuron.2017.06.041.

Bandettini, Peter A., Laurentius (Renzo) Huber, and Emily S. Finn. 2021. “Challenges and Opportunities of Mesoscopic Brain Mapping with FMRI.” Current Opinion in Behavioral Sciences 40:189–200. doi: 10.1016/j.cobeha.2021.06.002.

Beckett, Alexander JS, An T. Vu, Sinyeob Ahn, Salvatore Torrisi, Jonathan R. Polimeni, Essa Yacoub, Kawin Setsampop, Berkin Bilgic, Shajan Gunamony, Andreas Potthast, Peter Dietz, Yulin Chang, and David A. Feinberg. 2022. “Evaluation of Single-Shot EPI with Sub-Millimeter Resolution FMRI on the Next-Generation 7T Brain Scanner.”

Birn, Rasmus M., Erin K. Molloy, Rémi Patriat, Taurean Parker, Timothy B. Meier, Gregory R. Kirk, Veena A. Nair, M. Elizabeth Meyerand, and Vivek Prabhakaran. 2013. “The Effect of Scan Length on the Reliability of Resting-State FMRI Connectivity Estimates.” NeuroImage 83:550–58. doi: 10.1016/j.neuroimage.2013.05.099.

Biswal, Bharat, F. Zerrin Yetkin, Victor M. Haughton, and James S. Hyde. 1995. “Functional Connectivity in the Motor Cortex of Resting Human Brain Using Echo-Planar Mri.” Magnetic Resonance in Medicine 34(4):537–41. doi: 10.1002/mrm.1910340409.

Bollmann, Saskia, and Markus Barth. 2021. “New Acquisition Techniques and Their Prospects for the Achievable Resolution of FMRI.” Progress in Neurobiology 207:101936. doi: 10.1016/j.pneurobio.2020.101936.

Breuer, Felix A., Martin Blaimer, Matthias F. Mueller, Nicole Seiberlich, Robin M. Heidemann, Mark A. Griswold, and Peter M. Jakob. 2006. “Controlled Aliasing in Volumetric Parallel Imaging (2D CAIPIRINHA).” Magnetic Resonance in Medicine 55(3):549–56. doi: 10.1002/mrm.20787.

Chai, Yuhui, Linqing Li, Yicun Wang, Laurentius (Renzo) Huber, Benedikt A. Poser, Jeff Duyn, and Peter A. Bandettini. 2021. “Magnetization Transfer Weighted EPI Facilitates Cortical Depth Determination in Native FMRI Space.” NeuroImage 242:118455. doi: 10.1016/j.neuroimage.2021.118455.

Chang, Wei-Tang, Stephanie Langella, Min Sung Seo, Khoi Huynh, Pew-Thian Yap, Weili Lin, and Kelly Giovanello1. 2022. Cross-Layer Balance of Visuo-Hippocampal Functional Connectivity Is Associated With Episodic Memory Recognition Accuracy. preprint. In Review. doi: 10.21203/rs.3.rs-1789565/v1.

Chen, J. Jean, and Claudine J. Gauthier. 2021. “The Role of Cerebrovascular-Reactivity Mapping in Functional MRI: Calibrated FMRI and Resting-State FMRI.” Frontiers in Physiology 12:657362. doi: 10.3389/fphys.2021.657362.

Cordes, D., V. M. Haughton, K. Arfanakis, G. J. Wendt, P. A. Turski, C. H. Moritz, M. A. Quigley, and M. E. Meyerand. 2000. “Mapping Functionally Related Regions of Brain with Functional Connectivity MR Imaging.” AJNR. American Journal of Neuroradiology 21(9):1636–44.

De Luca, M., C. F. Beckmann, N. De Stefano, P. M. Matthews, and S. M. Smith. 2006. “FMRI Resting State Networks Define Distinct Modes of Long-Distance Interactions in the Human Brain.” NeuroImage 29(4):1359–67. doi: 10.1016/j.neuroimage.2005.08.035.

Deshpande, Gopikrishna, Yun Wang, and Jennifer Robinson. 2022. “Resting State FMRI Connectivity Is Sensitive to Laminar Connectional Architecture in the Human Brain.” Brain Informatics 9(1):2. doi: 10.1186/s40708-021-00150-4.

Deshpande, Gopikrishna, Xinyu Zhao, and Jennifer Robinson. 2022. “Functional Parcellation of the Hippocampus Based on Its Layer-Specific Connectivity with Default Mode and Dorsal Attention Networks.” NeuroImage 254:119078. doi: 10.1016/j.neuroimage.2022.119078.

Dworkin, Jordan D., Kristin A. Linn, Erin G. Teich, Perry Zurn, Russell T. Shinohara, and Danielle S. Bassett. 2020. “The Extent and Drivers of Gender Imbalance in Neuroscience Reference Lists.” Nature Neuroscience 23(8):918–26. doi: 10.1038/s41593-020-0658-y.

Feinberg, David A., Salvatore Torrisi, Alexander JS Beckett, Rüdiger Stirnberg, Tony Stöcker, Philipp Ehses, and Laurentius (Renzo) Huber. 2022. “Sub-0.1 Microliter CBV FMRI on the Next Generation 7T Scanner.”

Felleman, Daniel J., and David C. Van Essen. 1991. “Distributed Hierarchical Processing in the Primate Cerebral Cortex.” Cerebral Cortex 1(1):1–47. doi: 10.1093/cercor/1.1.1-a.

Finn, Emily S., and Peter A. Bandettini. 2021. “Movie-Watching Outperforms Rest for Functional Connectivity-Based Prediction of Behavior.” NeuroImage 235:117963. doi: 10.1016/j.neuroimage.2021.117963.

Finn, Emily S., Laurentius (Renzo) Huber, and Peter A. Bandettini. 2021. “Higher and Deeper: Bringing Layer FMRI to Association Cortex.” Progress in Neurobiology 207:101930. doi: 10.1016/j.pneurobio.2020.101930.

Finn, Emily S., Laurentius (Renzo) Huber, David C. Jangraw, Peter J. Molfese, and Peter A. Bandettini. 2019. “Layer-Dependent Activity in Human Prefrontal Cortex during Working Memory.” Nature Neuroscience 22(10):1687–95. doi: 10.1038/s41593-019-0487-z.

Fransson, Peter, and Guillaume Marrelec. 2008. “The Precuneus/Posterior Cingulate Cortex Plays a Pivotal Role in the Default Mode Network: Evidence from a Partial Correlation Network Analysis.” NeuroImage 42(3):1178–84. doi: 10.1016/j.neuroimage.2008.05.059.

Gau, Rémi, Stephanie Noble, Katja Heuer, Katherine L. Bottenhorn, Isil P. Bilgin, Yu-Fang Yang, Julia M. Huntenburg, Johanna M. M. Bayer, et al., 2021. “Brainhack: Developing a Culture of Open, Inclusive, Community-Driven Neuroscience.” Neuron 109(11):1769–75. doi: 10.1016/j.neuron.2021.04.001.

Glasser, Matthew F., Timothy S. Coalson, Emma C. Robinson, Carl D. Hacker, John Harwell, Essa Yacoub, Kamil Ugurbil, Jesper Andersson, Christian F. Beckmann, Mark Jenkinson, Stephen M. Smith, and David C. Van Essen. 2016. “A Multi-Modal Parcellation of Human Cerebral Cortex.” Nature 536(7615):171–78. doi: 10.1038/nature18933.

Godlove, D. C., A. Maier, G. F. Woodman, and J. D. Schall. 2014. “Microcircuitry of Agranular Frontal Cortex: Testing the Generality of the Canonical Cortical Microcircuit.” Journal of Neuroscience 34(15):5355–69. doi: 10.1523/JNEUROSCI.5127-13.2014.

Goense, Jozien, Yvette Bohraus, and Nikos K. Logothetis. 2016. “FMRI at High Spatial Resolution: Implications for BOLD-Models.” Frontiers in Computational Neuroscience 10. doi: 10.3389/fncom.2016.00066.

Gorgolewski, Krzysztof J., Tibor Auer, Vince D. Calhoun, R. Cameron Craddock, Samir Das, Eugene P. Duff, Guillaume Flandin, Satrajit S. Ghosh, Tristan Glatard, Yaroslav O. Halchenko, Daniel A. Handwerker, Michael Hanke, David Keator, Xiangrui Li, Zachary Michael, Camille Maumet, B. Nolan Nichols, Thomas E. Nichols, John Pellman, Jean-Baptiste Poline, Ariel Rokem, Gunnar Schaefer, Vanessa Sochat, William Triplett, Jessica A. Turner, Gaël Varoquaux, and Russell A. Poldrack. 2016. “The Brain Imaging Data Structure, a Format for Organizing and Describing Outputs of Neuroimaging Experiments.” Scientific Data 3(1):160044. doi: 10.1038/sdata.2016.44.

Griswold, Mark A., Peter M. Jakob, Robin M. Heidemann, Mathias Nittka, Vladimir Jellus, Jianmin Wang, Berthold Kiefer, and Axel Haase. 2002. “Generalized Autocalibrating Partially Parallel Acquisitions (GRAPPA).” Magnetic Resonance in Medicine 47(6):1202–10. doi: 10.1002/mrm.10171.

Guidi, Maria, Laurentius (Renzo) Huber, Leonie Lampe, Alberto Merola, Kristin Ihle, and Harald E. Möller. 2020. “Cortical Laminar Resting-state Signal Fluctuations Scale with the Hypercapnic Blood Oxygenation Level-dependent Response.” Human Brain Mapping 41(8):2014–27. doi: 10.1002/hbm.24926.

Haarsma, Joost, Peter Kok, and Michael Browning. 2020. The Promise of Layer-Specific Neuroimaging for Testing Predictive Coding Theories of Psychosis. preprint. PsyArXiv. doi: 10.31234/osf.io/xeyv7.

Havlicek, Martin, Alard Roebroeck, Karl Friston, Anna Gardumi, Dimo Ivanov, and Kamil Uludag. 2015. “Physiologically Informed Dynamic Causal Modeling of FMRI Data.” NeuroImage 122:355–72. doi: 10.1016/j.neuroimage.2015.07.078.

He, B. J. 2011. “Scale-Free Properties of the Functional Magnetic Resonance Imaging Signal during Rest and Task.” Journal of Neuroscience 31(39):13786–95. doi: 10.1523/JNEUROSCI.2111-11.2011.

Heid, Oliver. 1997. “Method for the Phase Correction of Nuclear Magnetic Resonance Signals.”

Heinzle, Jakob, Peter J. Koopmans, Hanneke E. M. den Ouden, Sudhir Raman, and Klaas Enno Stephan. 2016. “A Hemodynamic Model for Layered BOLD Signals.” NeuroImage 125:556–70. doi: 10.1016/j.neuroimage.2015.10.025.

Hua, Jun, Craig K. Jones, Qin Qin, and Peter C. M. van Zijl. 2013. “Implementation of Vascular-Space-Occupancy MRI at 7T: 3D MT-VASO MRI at 7T.” Magnetic Resonance in Medicine 69(4):1003–13. doi: 10.1002/mrm.24334.

Huber, Laurentius (Renzo), Emily S. Finn, Yuhui Chai, Rainer Goebel, Rüdiger Stirnberg, Tony Stöcker, Sean Marrett, Kamil Uludag, Seong-Gi Kim, SoHyun Han, Peter A. Bandettini, and Benedikt A. Poser. 2021. “Layer-Dependent Functional Connectivity Methods.” Progress in Neurobiology 207:101835. doi: 10.1016/j.pneurobio.2020.101835.

Huber, Laurentius (Renzo), Emily S. Finn, Daniel A. Handwerker, Marlene Bönstrup, Daniel R. Glen, Sriranga Kashyap, Dimo Ivanov, Natalia Petridou, Sean Marrett, Jozien Goense, Benedikt A. Poser, and Peter A. Bandettini. 2020. “Sub-Millimeter FMRI Reveals Multiple Topographical Digit Representations That Form Action Maps in Human Motor Cortex.” NeuroImage 208:116463. doi: 10.1016/j.neuroimage.2019.116463.

Huber, Laurentius (Renzo), Daniel A. Handwerker, David C. Jangraw, Gang Chen, Andrew Hall, Carsten Stüber, Javier Gonzalez-Castillo, Dimo Ivanov, Sean Marrett, Maria Guidi, Jozien Goense, Benedikt A. Poser, and Peter A. Bandettini. 2017. “High-Resolution CBV-FMRI Allows Mapping of Laminar Activity and Connectivity of Cortical Input and Output in Human M1.” Neuron 96(6):1253-1263.e7. doi: 10.1016/j.neuron.2017.11.005.

Huber, Laurentius (Renzo), Dimo Ivanov, Daniel A. Handwerker, Sean Marrett, Maria Guidi, Kâmil Uludağ, Peter A. Bandettini, and Benedikt A. Poser. 2018. “Techniques for Blood Volume FMRI with VASO: From Low-Resolution Mapping towards Sub-Millimeter Layer-Dependent Applications.” NeuroImage 164:131–43. doi: 10.1016/j.neuroimage.2016.11.039.

Huber, Laurentius (Renzo), Lisa Kronbichler, Ruediger Stirnberg, Philipp Ehses, Tony Stoecker, Sara Fernandez-Cabello, Benedikt A. Poser, and Martin Kronbichler. 2022. Evaluating the Capabilities and Challenges of Layer-FMRI VASO at 3T. preprint. Neuroscience. doi: 10.1101/2022.07.26.501554.

Huber, Laurentius (Renzo), Benedikt A. Poser, Peter A. Bandettini, Kabir Arora, Konrad Wagstyl, Shinho Cho, Jozien Goense, Nils Nothnagel, Andrew Tyler Morgan, Job van den Hurk, Anna K. Müller, Richard C. Reynolds, Daniel R. Glen, Rainer Goebel, and Omer Faruk Gulban. 2021. “LayNii: A Software Suite for Layer-FMRI.” NeuroImage 237:118091. doi: 10.1016/j.neuroimage.2021.118091.

Huber, Laurentius (Renzo), Benedikt A. Poser, Amanda L. Kaas, Elizabeth J. Fear, Sebastian Desbach, Jason Berwick, Rainer Goebel, Robert Turner, and Aneurin J. Kennerley. 2020. Validating Layer-Specific VASO across Species. preprint. Neuroscience. doi: 10.1101/2020.07.24.219378.

Huotari, Niko, Lauri Raitamaa, Heta Helakari, Janne Kananen, Ville Raatikainen, Aleksi Rasila, Timo Tuovinen, Jussi Kantola, Viola Borchardt, Vesa J. Kiviniemi, and Vesa O. Korhonen. 2019. “Sampling Rate Effects on Resting State FMRI Metrics.” Frontiers in Neuroscience 13:279. doi: 10.3389/fnins.2019.00279.

Jia, Ke, Elisa Zamboni, Valentin Kemper, Catarina Rua, Nuno Reis Goncalves, Adrian Ka Tsun Ng, Christopher T. Rodgers, Guy Williams, Rainer Goebel, and Zoe Kourtzi. 2020. “Recurrent Processing Drives Perceptual Plasticity.” Current Biology 30(21):4177-4187.e4. doi: 10.1016/j.cub.2020.08.016.

Kazan, Samira M., Laurentius (Renzo) Huber, Guillaume Flandin, Dimo Ivanov, Peter Bandettini, and Nikolaus Weiskopf. 2017. “Physiological Basis of Vascular Autocalibration (VasA): Comparison to Hypercapnia Calibration Methods: BOLD Signal Calibration with Low-Frequency Fluctuations.” Magnetic Resonance in Medicine 78(3):1168–73. doi: 10.1002/mrm.26494.

Kazan, Samira M., Siawoosh Mohammadi, Martina F. Callaghan, Guillaume Flandin, Laurentius (Renzo) Huber, Robert Leech, Aneurin Kennerley, Christian Windischberger, and Nikolaus Weiskopf. 2016. “Vascular Autorescaling of FMRI (VasA FMRI) Improves Sensitivity of Population Studies: A Pilot Study.” NeuroImage 124:794–805. doi: 10.1016/j.neuroimage.2015.09.033.

Kellman, Peter, and Elliot R. McVeigh. 2005. “Image Reconstruction in SNR Units: A General Method for SNR Measurement.” Magnetic Resonance in Medicine 54(6):1439–47. doi: 10.1002/mrm.20713.

Klein, Arno, and Jason Tourville. 2012. “101 Labeled Brain Images and a Consistent Human Cortical Labeling Protocol.” Frontiers in Neuroscience 6. doi: 10.3389/fnins.2012.00171.

Klein, Richard, Tyler Morgan, Burak Akin, Laurentius (Renzo) Huber, and Peter A. Bandettini. 2022. “Laminar-Dependent Functional Connectivity between Area V4 and Frontopariental Cortex.”

Knapen, Tomas. 2021. “Topographic Connectivity Reveals Task-Dependent Retinotopic Processing throughout the Human Brain.” Proceedings of the National Academy of Sciences 118(2):e2017032118. doi: 10.1073/pnas.2017032118.

Kriegeskorte, Nikolaus. 2008. “Representational Similarity Analysis – Connecting the Branches of Systems Neuroscience.” Frontiers in Systems Neuroscience. doi: 10.3389/neuro.06.004.2008.

Lauren, Peter, Daniel R. Glen, Richard C. Reynolds, Salvatore Torrisi, and Paul A. Taylor. 2022. “Using Clipping Planes to Analyze Brain Data in Suma.”

Lewis, Laura D., Kawin Setsompop, Bruce R. Rosen, and Jonathan R. Polimeni. 2016. “Fast FMRI Can Detect Oscillatory Neural Activity in Humans.” Proceedings of the National Academy of Sciences 113(43). doi: 10.1073/pnas.1608117113.

Li, Xiangrui, Paul S. Morgan, John Ashburner, Jolinda Smith, and Christopher Rorden. 2016. “The First Step for Neuroimaging Data Analysis: DICOM to NIfTI Conversion.” Journal of Neuroscience Methods 264:47–56. doi: 10.1016/j.jneumeth.2016.03.001.

Liu, Peiying, Andrew C. Hebrank, Karen M. Rodrigue, Kristen M. Kennedy, Denise C. Park, and Hanzhang Lu. 2013. “A Comparison of Physiologic Modulators of FMRI Signals: Comparison of Physiologic Modulators of FMRI.” Human Brain Mapping 34(9):2078–88. doi: 10.1002/hbm.22053.

Lowe, M. J., B. J. Mock, and J. A. Sorenson. 1998. “Functional Connectivity in Single and Multislice Echoplanar Imaging Using Resting-State Fluctuations.” NeuroImage 7(2):119–32. doi: 10.1006/nimg.1997.0315.

Lu, Hanzhang, Xavier Golay, James J. Pekar, and Peter C. M. van Zijl. 2003. “Functional Magnetic Resonance Imaging Based on Changes in Vascular Space Occupancy.” Magnetic Resonance in Medicine 50(2):263–74. doi: 10.1002/mrm.10519.

Lu, Hanzhang, Peter C. M. van Zijl, Jeroen Hendrikse, and Xavier Golay. 2004. “Multiple Acquisitions with Global Inversion Cycling (MAGIC): A Multislice Technique for Vascular-Space-Occupancy Dependent FMRI.” Magnetic Resonance in Medicine 51(1):9–15. doi: 10.1002/mrm.10659.

Ma, Ying, Mohammed A. Shaik, Mariel G. Kozberg, Sharon H. Kim, Jacob P. Portes, Dmitriy Timerman, and Elizabeth M. C. Hillman. 2016. “Resting-State Hemodynamics Are Spatiotemporally Coupled to Synchronized and Symmetric Neural Activity in Excitatory Neurons.” Proceedings of the National Academy of Sciences 113(52). doi: 10.1073/pnas.1525369113.

Mandelkow, H., J. A. de Zwart, and J. H. Duyn. 2017. “Effects of Spatial FMRI Resolution on the Classification of Naturalistic Movies.” NeuroImage 162:45–55. doi: 10.1016/j.neuroimage.2017.08.053.

Markiewicz, Christopher J., Krzysztof J. Gorgolewski, Franklin Feingold, Ross Blair, Yaroslav O. Halchenko, Eric Miller, Nell Hardcastle, Joe Wexler, Oscar Esteban, Mathias Goncavles, Anita Jwa, and Russell Poldrack. 2021. “The OpenNeuro Resource for Sharing of Neuroscience Data.” ELife 10:e71774. doi: 10.7554/eLife.71774.

Markov, Nikola T., and Henry Kennedy. 2013. “The Importance of Being Hierarchical.” Current Opinion in Neurobiology 23(2):187–94. doi: 10.1016/j.conb.2012.12.008.

Markuerkiaga, Irati, Markus Barth, and David G. Norris. 2016. “A Cortical Vascular Model for Examining the Specificity of the Laminar BOLD Signal.” NeuroImage 132:491–98. doi: 10.1016/j.neuroimage.2016.02.073.

McColgan, Peter, Julie Joubert, Sarah J. Tabrizi, and Geraint Rees. 2020. “The Human Motor Cortex Microcircuit: Insights for Neurodegenerative Disease.” Nature Reviews Neuroscience 21(8):401–15. doi: 10.1038/s41583-020-0315-1.

Menon, Ravi S., and Bradley G. Goodyear. 1999. “Submillimeter Functional Localization in Human Striate Cortex Using BOLD Contrast at 4 Tesla: Implications for the Vascular Point-Spread Function.” Magnetic Resonance in Medicine 41(2):230–35. doi: 10.1002/(SICI)1522-2594(199902)41:2<230::AID-MRM3>3.0.CO;2-O.

van Mourik, Tim, Peter J. Koopmans, Lauren J. Bains, David G. Norris, and Janneke F. M. Jehee. 2021. Investigation of Layer Specific BOLD in the Human Visual Cortex during Visual Attention. preprint. Neuroscience. doi: 10.1101/2021.02.07.430129.

Müller, Anna Katharina, Heynckes, Miriam, Wiggins, J, Christopher, Gulban, Faruk, Omer, Chai, Yuhui, Benedikt A. Poser, and Laurentius (Renzo) Huber. 2021. “Whole Brain Layer-FMRI: An Open Dataset for Methods Benchmarking.”

Müller, Anna Katharina, and Laurentius (Renzo) Huber. 2020. “Whole Brain Layer-FMRI Connectome: An Open Dataset.” doi: 10.5281/ZENODO.6860807.

Naselaris, Thomas, Kendrick N. Kay, Shinji Nishimoto, and Jack L. Gallant. 2011. “Encoding and Decoding in FMRI.” NeuroImage 56(2):400–410. doi: 10.1016/j.neuroimage.2010.07.073.

Pais-Roldán, Patricia, Seong Dae Yun, Nicola Palomero-Gallagher, and N. Jon Shah. 2020. Cortical Depth-Dependent Human FMRI of Resting-State Networks Using EPIK. preprint. Neuroscience. doi: 10.1101/2020.12.07.414144.

Polimeni, Jonathan R., M. Mianciardi, B. Keil, and L. L. Wald. 2010. “Cortical Depth Dependence of Physiological Fluctuations and Whole-Brain Resting-State Functional Connectivity at 7T.”

Polimeni, Jonathan R., Ville Renvall, Natalia Zaretskaya, and Bruce Fischl. 2018. “Analysis Strategies for High-Resolution UHF-FMRI Data.” NeuroImage 168:296–320. doi: 10.1016/j.neuroimage.2017.04.053.

Poser, B. A., P. J. Koopmans, T. Witzel, L. L. Wald, and M. Barth. 2010. “Three Dimensional Echo-Planar Imaging at 7 Tesla.” NeuroImage 51(1):261–66. doi: 10.1016/j.neuroimage.2010.01.108.

Poser, Benedikt A., Demo Ivanov, Kemper, VG, Kannengiesser, SA, Uludag, K, and Barth, M. 2013. “CAIPIRINHA-Accelerated 3D EPI for High Temporal and / or Spatial Resolution EPI Acquisitions.”

Schluppeck, Denis, Rosa-Maria Sanchez-Panchuelo, and Susan T. Francis. 2018. “Exploring Structure and Function of Sensory Cortex with 7 T MRI.” NeuroImage 164:10–17. doi: 10.1016/j.neuroimage.2017.01.081.

Schneider, Marian, Valentin G. Kemper, Thomas C. Emmerling, Federico De Martino, and Rainer Goebel. 2019. “Columnar Clusters in the Human Motion Complex Reflect Consciously Perceived Motion Axis.” Proceedings of the National Academy of Sciences 116(11):5096–5101. doi: 10.1073/pnas.1814504116.

Scouten, A., and R. T. Constable. 2007. “Applications and Limitations of Whole-Brain MAGIC VASO Functional Imaging.” Magnetic Resonance in Medicine 58(2):306–15. doi: 10.1002/mrm.21273.

Self, Matthew W., Timo van Kerkoerle, Rainer Goebel, and Pieter R. Roelfsema. 2019. “Benchmarking Laminar FMRI: Neuronal Spiking and Synaptic Activity during Top-down and Bottom-up Processing in the Different Layers of Cortex.” NeuroImage 197:806–17. doi: 10.1016/j.neuroimage.2017.06.045.

Sharoh, Daniel, Tim van Mourik, Lauren J. Bains, Katrien Segaert, Kirsten Weber, Peter Hagoort, and David G. Norris. 2019. “Laminar Specific FMRI Reveals Directed Interactions in Distributed Networks during Language Processing.” Proceedings of the National Academy of Sciences 116(42):21185–90. doi: 10.1073/pnas.1907858116.

Sotero, Roberto C., Aleksandra Bortel, Shmuel Naaman, Victor M. Mocanu, Pascal Kropf, Martin Villeneuve, and Amir Shmuel. 2015. “Laminar Distribution of Phase-Amplitude Coupling of Spontaneous Current Sources and Sinks.” Frontiers in Neuroscience 9. doi: 10.3389/fnins.2015.00454.

Stephan, K. E., F. H. Petzschner, L. Kasper, J. Bayer, K. V. Wellstein, G. Stefanics, K. P. Pruessmann, and J. Heinzle. 2019. “Laminar FMRI and Computational Theories of Brain Function.” NeuroImage 197:699–706. doi: 10.1016/j.neuroimage.2017.11.001.

Stirnberg, Rüdiger, and Tony Stöcker. 2021. “Segmented K-space Blipped-controlled Aliasing in Parallel Imaging for High Spatiotemporal Resolution EPI.” Magnetic Resonance in Medicine 85(3):1540–51. doi: 10.1002/mrm.28486.

Torrisi, Salvatore, Peter Lauren, Paul A. Taylor, Suhyung Park, David A. Feinberg, and Daniel R. Glen. 2021. “Creating Layered Surfaces to Visualize with AFNI + SUMA, with Applications to Laminar FMRI.”

Vanderwal, Tamara, Clare Kelly, Jeffrey Eilbott, Linda C. Mayes, and F. Xavier Castellanos. 2015. “Inscapes : A Movie Paradigm to Improve Compliance in Functional Magnetic Resonance Imaging.” NeuroImage 122:222–32. doi: 10.1016/j.neuroimage.2015.07.069.

Vizioli, Luca, Steen Moeller, Logan Dowdle, Mehmet Akçakaya, Federico De Martino, Essa Yacoub, and Kamil Uğurbil. 2021. “Lowering the Thermal Noise Barrier in Functional Brain Mapping with Magnetic Resonance Imaging.” Nature Communications 12(1):5181. doi: 10.1038/s41467-021-25431-8.

Webb, A. G. 2011. “Dielectric Materials in Magnetic Resonance.” Concepts in Magnetic Resonance Part A 38A(4):148–84. doi: 10.1002/cmr.a.20219.

Weldon, Kimberly B., and Cheryl A. Olman. 2021. “Forging a Path to Mesoscopic Imaging Success with Ultra-High Field Functional Magnetic Resonance Imaging.” Philosophical Transactions of the Royal Society B: Biological Sciences 376(1815):20200040. doi: 10.1098/rstb.2020.0040.

Wu, Shian, Yi Liu, Yonggang Zheng, Jixin Dong, and Duojia Pan. 2008. “The TEAD/TEF Family Protein Scalloped Mediates Transcriptional Output of the Hippo Growth-Regulatory Pathway.” Developmental Cell 14(3):388–98. doi: 10.1016/j.devcel.2008.01.007.

Yang, Jiajia, Laurentius (Renzo) Huber, Yinghua Yu, and Peter A. Bandettini. 2021. “Linking Cortical Circuit Models to Human Cognition with Laminar FMRI.” Neuroscience & Biobehavioral Reviews 128:467–78. doi: 10.1016/j.neubiorev.2021.07.005.

Yun, Seong Dae, Patricia Pais-Roldán, Nicola Palomero-Gallagher, and N. Jon Shah. 2022. “Mapping of Whole-cerebrum Resting-state Networks Using Ultra-high Resolution Acquisition Protocols.” Human Brain Mapping 43(11):3386–3403. doi: 10.1002/hbm.25855.

Yushkevich, Paul A., Joseph Piven, Heather Cody Hazlett, Rachel Gimpel Smith, Sean Ho, James C. Gee, and Guido Gerig. 2006. “User-Guided 3D Active Contour Segmentation of Anatomical Structures: Significantly Improved Efficiency and Reliability.” NeuroImage 31(3):1116–28. doi: 10.1016/j.neuroimage.2006.01.015.

